# Cholecystokinin-expressing (CCK+) basket cells are key controllers of theta-gamma coupled rhythms in the hippocampus

**DOI:** 10.1101/2022.06.01.494440

**Authors:** Alexandra P Chatzikalymniou, Spandan Sengupta, Jeremie Lefebvre, Frances K Skinner

## Abstract

Various inhibitory cell types underlie cognitively important brain rhythms and their couplings. Precisely how this is manifest is unclear. Complex network interactions make an understanding of inhibitory cell contributions extremely difficult and using experiments alone is insufficient. Using detailed biophysical models, we obtain hypotheses of how theta and gamma rhythms in the hippocampus are generated and coupled. We find critical contributions by parvalbumin-expressing (PV+) basket cells (BCs), cholecystokinin-expressing (CCK+) BCs and bistratified cells. Based on this, we develop and explore a population rate model and predict that CCK+BCs exert more control relative to PV+BCs for theta-gamma coupling, and that theta frequencies are more strongly affected by PV+BC to CCK+BC coupling relative to CCK+BC to PV+BC. As specific inhibitory cell types can be targeted during behaviour, it is possible to test these predictions. Our work shows that combining models at different scales creates new insights that otherwise would not be revealed.

## Introduction

We are now firmly in an era of distinct interneuron or inhibitory cell types, and it is apparent that the different interneuron types need to be considered from functional viewpoints in cognition and behaviour (***Fishell and Kepecs, 2020***; ***Kepecs and Fishell, 2014***; ***Klausberger and Somogyi, 2008***; ***McBain and Fisahn, 2001***). There is a wide range of rhythmic brain frequencies that have been recorded in local field potentials (LFPs) and electroencephalograms (EEGs) arising from dynamic interactions of excitatory and inhibitory cell types in brain circuits (***Buzsáki, 2006***). Moreover, it is becoming clear that cross-frequency coupling (CFC) between high and low frequency rhythms may play functional roles in sensory, motor and cognitive events (***Canolty and Knight, 2010***).

The hippocampus is a heavily studied brain structure that expresses rhythmic activities with well-defined behavioural correlates (***Buzsaki, 2011***; ***Colgin, 2016***). In particular, the theta oscillation (≈3-12 Hz) is a prominent LFP rhythm that is most robustly recorded from the CA1 region of the hippocampus (***Buzsáki, 2002***). This rhythm enables multiple timescale organization of neuronal assemblies facilitating the spatio-temporal encoding of events during episodic recall (***Buzsáki and Moser, 2013***). Indeed, it has been suggested that ‘the single theta cycle is a functional unit capable of representing distinct temporal-spatial content at different phases’ (***Wilson et al., 2015***). Interestingly, disruptions of hippocampal theta rhythms are associated with memory impairments (***Robbe and Buzsáki, 2009***). Higher frequency gamma rhythms (≈ 20-100 Hz) occur nested in these theta oscillations (***Buzsáki et al., 1983***; ***Colgin, 2015***), and CFC of these rhythms are considered central to hippocampal function (***Colgin et al., 2009***; ***Tort et al., 2008***) with possible causal roles (***Radiske et al., 2020***). Dynamic modulation of theta/gamma rhythms during sleep (***Bandarabadi et al., 2019***), visual exploration (***Kragel et al., 2020***) and association with working memory (***Axmacher et al., 2010***; ***Lega et al., 2016***) have been documented. It is perhaps not surprising that mounting evidence points to specific changes in theta-gamma coupling with memory impairments, disease and its progression (***Goutagny et al., 2013***; ***Hamm et al., 2015***; ***Karlsson et al., 2022***; ***Kitchigina, 2018***; ***Musaeus et al., 2020***; ***Zhang et al., 2016***).

While it has been shown that specific cell types contribute in particular ways to generating theta and gamma rhythms, exactly how they contribute to control these rhythms and how they mutually interact is far from clear. In trying to understand the contributions of specific cell types, it has long been noted that there is a separation between perisomatically and dendritically targeting interneuron types onto pyramidal cells (***Freund and Buzsáki, 1996***; ***McBain and Fisahn, 2001***). Thus, one can consider a dichotomy in the inhibitory control of pyramidal cell excitability. Further, ***Freund*** (***2003***) has described another dichotomy of perisomatically targeting cholecystokinin-expressing (CCK+) and parvalbumin-expressing (PV+) basket cells (BCs) in a ‘rhythm and mood’ fashion (***Freund, 2003***; ***Freund and Katona, 2007***), with PV+BCs contributing in a precise clockwork fashion and CCK+BCs in a highly modulatory way. Many mathematical models have been created to help decipher the complexity of excitatory and inhibitory interactions underlying theta and gamma rhythms and their couplings (***Ferguson and Skinner, 2015***). In particular, circuit models that include explicitly-identified and characterized cell types have been developed (***Bezaire et al., 2016b***; ***Ferguson et al., 2017***; ***Ecker et al., 2020***).

Bezaire and colleagues developed a full-scale model (FSM) of the CA1 hippocampus that exhibits theta and gamma rhythms (***Bezaire et al., 2016b***). This is a biophysically detailed microcircuit model with 338,740 cells that includes pyramidal (PYR) cells, PV+BCs, axo-axonic cells (AACs), bistratified cells (BiCs), CCK+BCs, Schaeffer Collateral-associated (SCA) cells, oriens lacunosum-moleculare (OLM) cells, neurogliaform (NGF) cells, and ivy cells. The FSM provides a realistic representation of the hippocampus which is grounded upon a previously compiled, extensive quantitative analysis (***Bezaire and Soltesz, 2013***). The authors used their FSM to shed light on the generation mechanism of theta rhythms by describing the activities of the nine cell types. In broad terms, the FSM distinguished the importance of certain cell types against others, and predicted that cell type variability is necessary for theta rhythms to occur.

The very complexity of the FSM poses a challenge in the elucidation of explicit mechanisms producing theta rhythms and theta-gamma coupling from an inhibitory cell type perspective. On the one hand, we very much need cellular-based network models to help us untangle the various facets of the interacting dynamics of the network system that produces theta and gamma rhythms and their coupling as this cannot come from experiments alone. Simply identifying and targeting the many different inhibitory cell types during ongoing behaviours and rhythmic activities is an immense, ongoing experimental challenge that is extremely laborious (***Dudok et al., 2021a***,b; ***Leão et al., 2012***). On the other hand, the presence of nonlinearity, high-dimensionality and degeneracy in detailed models makes it difficult to extract explicit mechanisms.

In this paper, we build on our previous work (***Chatzikalymniou et al., 2021***) and take a deep dive into the FSM of ***Bezaire et al***. (***2016b***). We leverage these examinations to develop precise hypotheses of how theta and gamma rhythms are generated, interact with each other, and how the different cell types mediate these interactions. We identify four cell types (PYR cells, PV+BCs, BiCs, CCK+BCs) that are deemed essential, and we confirm this by selective deactivation of their interconnections. We build a population rate model (PRM) using these four cell types and find a parameter set that can capture the developed hypotheses. We then do systematic parameter explorations of the PRM to expose dynamic ‘balances’ underlying the rhythmic expressions in the network system. We exploit detailed visualizations of the system and predict that CCK+BCs exhibit a much higher degree of control relative to PV+BCs for the dominance of either theta or gamma rhythms in the network system, and thus theta-gamma coupling. Given that it is possible to specifically target PV+BCs and CCK+BCs in the behaving animal (***Dudok et al., 2021a***), this prediction opens up a plethora of possibilities that can be experimentally examined. Moreover, our work shows that developing, analyzing and combining mathematical model types at different scales and abstraction brings about insights that would not be possible if only one model type were used. Interestingly, model diversity has recently been noted as needed to be able to have an immense impact on our understanding of the brain (***Eriksson et al., 2022***).

## Results

To have a starting basis for examinations using the FSM, we reproduced theta-gamma rhythmic output in the FSM of ***Bezaire et al***. (***2016b***). This is shown in FIGURE 1. In FIGURE 1A, we present a stylized schematic of the nine different cell types in the FSM. Connections between the different cell types are represented by grey lines between them. Grey circles by a given cell type indicates that they are interconnected, and incoming grey lines to the different cell types represent external drives. The thickness of the lines and circles is representative of the relative strength of these various connections. They reflect computed effective weights (EWs) and external drives in the FSM. The specific values are given in the Methods (see TABLE 1 for drives, and TABLES 2-3 for EWs). The raw unfiltered LFP output from the network model is shown in FIGURE 1B(i) with its power spectra in FIGURE 1B(ii). The raster plot output for each of the nine cell types is given in FIGURE 1C(i) for 4 seconds. As has been shown experimentally, these rhythms can be generated intra-hippocampally and do not require the medial septum (MS) (***Goutagny et al., 2009***; ***Jackson et al., 2011***). This does not mean that the MS is not important for theta rhythms, rather, it implies that the expression of these rhythms does not have explicit reliance on the MS and can be generated by hippocampal circuitry on its own.

**Figure 1.**
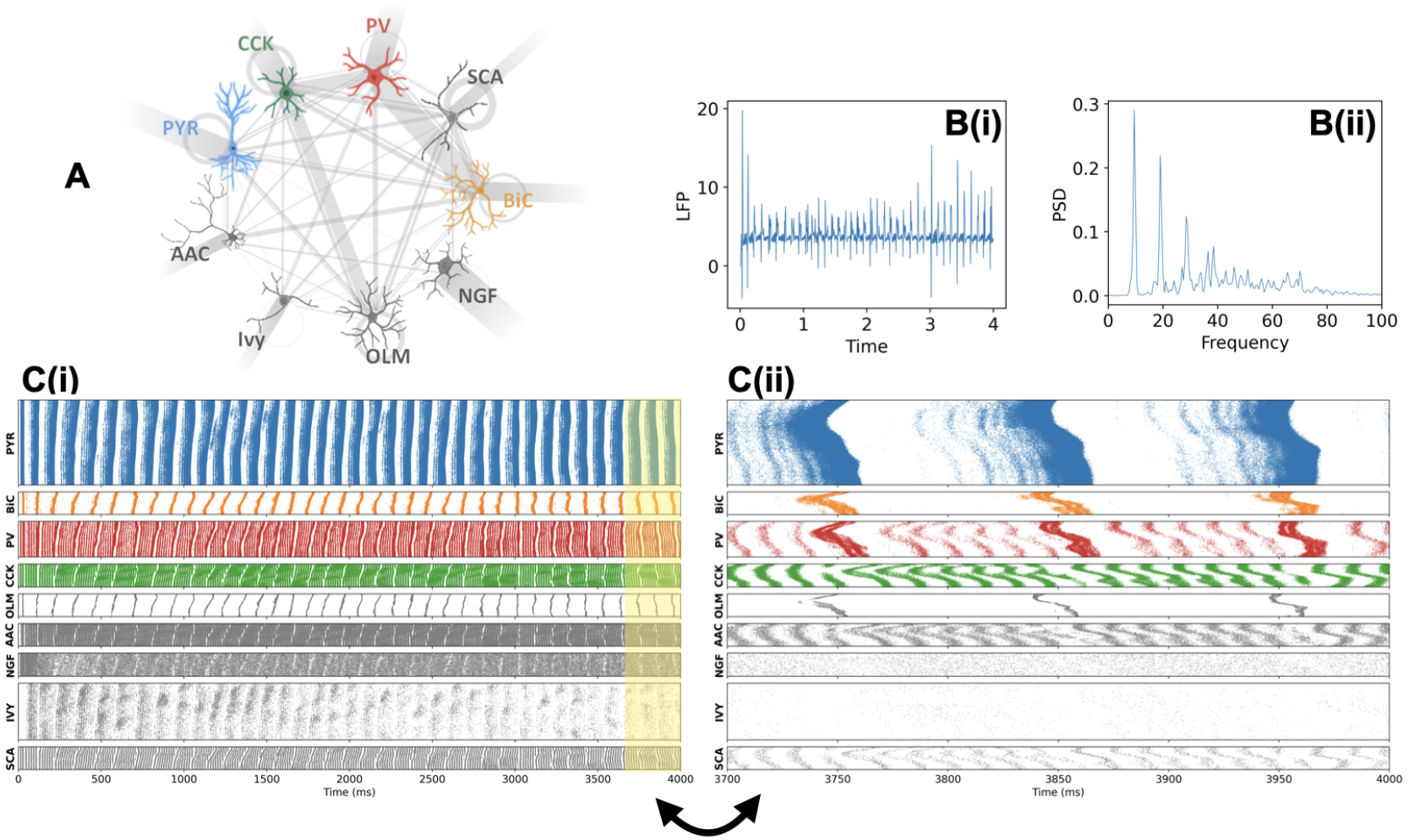
Output of the original full-scale model (FSM). **A**. Stylized schematic of the 9 different cell types that are present in the FSM. These cell types are pyramidal (PYR) cells, paravalbumin-expressing basket cells (PV+BCs), axo-axonic cells (AACs), bistratified cells (BiCs), cholecystokinin-expressing (CCK+) BCs, Schaeffer Collateral-associated (SCA) cells, oriens-lacunosum-moleculare (OLM) cells, neurogliaform (NGF) cells, and ivy cells. In the figure, they are referred to as PYR, PV, AAC, BiC, CCK, SCA, OLM, NGF, Ivy, respectively. Connectivities are represented by grey lines - see text for details. **B(i)** unfiltered LFP output (units of mV). **B(ii)** PSD (power spectral density) of LFP (units of mV^2^/Hz). **C(i)** 4 second (s) raster plot. **C(ii)** 300 millisecond (ms) raster plot, as taken from the last 300 ms of the simulation. Raster plots are of the excitatory cell types (PYR) and the eight inhibitory cell types (BiC, PV, CCK, OLM, AAC, NGF, IVY, SCA) as in the original CA1 FSM that includes 311,500 PYR (in blue); 2,210 BiC (in orange); 5,530 PV (in red); 3,600 CCK (in green); 1,640 OLM (in grey); 1,470 AAC (in grey); 3,580 NGF (in grey); 8,810 IVY (in grey) and 400 SCA (in grey). The colours match those of the stylized cells in the schematic of **A**.

**Table 1.** Number of cells and other input to each cell type in the FSM of ***Bezaire et al***. (***2016b***).

| Cell Type | Other Input | Number of Cells |
| --- | --- | --- |
| PYR cell | 2,914 | 311,500 |
| CCK-BC | 3,327 | 3,600 |
| PV-BC | 2,661 | 5,530 |
| BiC | 1,864 | 2,210 |
| OLM cell | 0 (NONE) | 1,640 |
| AAC | 1,117 | 1,470 |
| Ivy cell | 1,154 | 8,810 |
| SCA cell | 1,680 | 400 |
| NGF cell | 3,661 | 3,580 |

**Table 2.** Connectivities based on ***Bezaire et al***. (***2016b***) Appendix values for PYR cells, BiCs, CCK-BCs, PV-BCs and OLM cells.

| Connection Coupling | Number of Connections | Connection Probability | Synaptic Weight (nS) | EW Value (× 100) |
| --- | --- | --- | --- | --- |
| PYR->PYR | 197 | 0.00063 | 70.0 | 4.4 |
| PYR->BiC | 3 | 0.0014 | 5.7 | 0.80 |
| PYR->PV | 8 | 0.0015 | 2.1 | 0.32 |
| PYR->OLM | 13 | 0.0079 | 0.6 | 0.47 |
| PYR->CCK | - | - | - | 0 (NONE) |
| PYR->AAC | 1 | 0.00068 | 0.12 | 0.0082 |
| PYR->Ivy | - | - | - | 0 (NONE) |
| PYR->NGF | - | - | - | 0 (NONE) |
| PYR->SCA | - | - | - | 0 (NONE) |
| BiC->BiC | 16 | 0.0072 | 5.1 | 3.7 |
| BiC->PYR | 1410 | 0.0045 | 5.1 | 2.3 |
| BiC->CCK | 26 | 0.0072 | 8.0 | 5.8 |
| BiC->PV | 40 | 0.0072 | 90.0 | 64.8 |
| BiC->OLM | 29 | 0.018 | 0.2 | 0.35 |
| BiC->AAC | 11 | 0.0075 | 6.0 | 4.5 |
| BiC->Ivy | 12 | 0.0014 | 5.0 | 0.68 |
| BiC->NGF | - | - | - | 0 (NONE) |
| BiC->SCA | 3 | 0.0075 | 8.0 | 6.0 |
| CCK->CCK | 35 | 0.0097 | 3.6 | 3.5 |
| CCK->PV | 18 | 0.0033 | 72.0 | 23.8 |
| CCK->PYR | 1125 | 0.0036 | 4.2 | 1.5 |
| CCK->BiC | 7 | 0.0032 | 5.6 | 1.8 |
| CCK->OLM | 9 | 0.0055 | 5.6 | 3.1 |
| CCK->AAC | 5 | .0034 | 5.6 | 1.9 |
| CCK->Ivy | 20 | 0.0023 | 2.4 | 0.55 |
| CCK->NGF | - | - | - | 0 (NONE) |
| CCK->SCA | 3 | 0.0075 | 5.6 | 4.2 |
| PV->PV | 39 | 0.0071 | 1.6 | 1.1 |
| PV->PYR | 958 | 0.0031 | 2.2 | 0.7 |
| PV->CCK | 25 | 0.0069 | 1.2 | 0.8 |
| PV->BiC | 16 | 0.0072 | 2.9 | 2.1 |
| PV->OLM | - | - | - | 0 (NONE) |
| PV->AAC | 10 | 0.0068 | 0.12 | 0.8 |
| PV->Ivy | 13 | 0.0015 | 0.16 | 0.024 |
| PV->NGF | - | - | - | 0 (NONE) |
| PV->SCA | 2 | 0.005 | 0.6 | 0.3 |
| OLM->OLM | 6 | 0.0037 | 1.2 | 4.4 |
| OLM->PV | 27 | 0.0049 | 11.0 | 5.4 |
| OLM->CCK | 88 | 0.024 | 12.0 | 28.8 |
| OLM->BiC | 11 | 0.0050 | 1.1 | 0.55 |
| OLM->PYR | 1520 | 0.0049 | 3.0 | 1.5 |
| OLM->AAC | 7 | 0.0048 | 1.2 | 0.57 |
| OLM->Ivy | - | - | - | 0 (NONE) |
| OLM->NGF | 28 | 0.0078 | 0.98 | 0.77 |
| OLM->SCA | 10 | 0.025 | 1.5 | 3.8 |

**Table 3.** Connectivities based on ***Bezaire et al***. (***2016b***) Appendix values for AACs, Ivy cells, NGF cells and SCA cells.

| Connection Coupling | Number of Connections | Connection Probability | Synaptic Weight (nS) | EW Value (× 100) |
| --- | --- | --- | --- | --- |
| AAC->AAC | - | - | - | 0 (NONE) |
| AAC->PYR | 1271 | 0.0041 | 6.9 | 2.82 |
| AAC->BiC,PV,OLM,CCK,Ivy,NGF,SCA | - | - | - | 0 (NONE) |
| Ivy->Ivy | 24 | 0.0027 | 0.57 | 0.16 |
| Ivy->PYR | 1485 | 0.0048 | 0.41 | 0.20 |
| Ivy->BiC | 6 | 0.0027 | 0.77 | 0.21 |
| Ivy->PV | 15 | 0.0027 | 7.0 | 1.90 |
| Ivy->OLM | 25 | 0.015 | 0.57 | 0.87 |
| Ivy->CCK | 39 | 0.011 | 0.37 | 0.40 |
| Ivy->NGF | 11 | 0.0031 | 0.57 | 0.18 |
| Ivy->SCA | 5 | 0.013 | 0.37 | 0.46 |
| Ivy->AAC | 4 | 0.0027 | 0.57 | 0.16 |
| NGF->NGF | 17 | 0.0047 | 1.6 | 0.76 |
| NGF->PYR | 1218 | 0.0039 | 0.65 | 0.25 |
| NGF->BiC,PV,OLM,CCK,Ivy,SCA,AAC | - | - | - | 0 (NONE) |
| SCA->SCA | 6 | 0.015 | 6.0 | 9.0 |
| SCA->PYR | 1558 | 0.005 | 4.6 | 2.3 |
| SCA->BiC | 7 | 0.0032 | 3.6 | 1.15 |
| SCA->PV | 14 | 0.0025 | 7.8 | 1.95 |
| SCA->OLM | 8 | 0.0049 | 5.1 | 2.5 |
| SCA->CCK | 50 | 0.014 | 5.1 | 7.1 |
| SCA->Ivy | 44 | 0.0050 | 5.1 | 2.6 |
| SCA->NGF | - | - | - | 0 (NONE) |
| SCA->AAC | 4 | 0.0027 | 3.6 | 0.98 |

### Anatomy of a theta cycle in the full-scale model (FSM)

***Bezaire et al***. (***2016b***) identified a number of model properties as important for theta rhythms. They showed that diverse inhibitory cell types and the existence of recurrent excitation are necessary for the presence of these rhythms. They also performed mutation simulations and identified the cell types that were necessary for the rhythm versus those that could be removed without having much of an effect. Specifically SCA, OLM and ivy cell types were found to not be necessary. On the other hand, BiCs, PV+BCs and PYR cells were necessary and further, these cell types are strongly theta modulated (see Figure 5A in ***Bezaire et al***. (***2016b***), and we quantify this modulation in more detail in FIGURE 1-Supplement 1. The remaining cell types are CCK+BCs, AACs and NGF cells.

To further distill the essence of cell types that would be sufficient to have theta rhythms in the FSM, we examined external drives and connections between all of the cell types given their numbers and synaptic weights, where computed EWs are representative of cell numbers and synaptic weights (see Methods). Based on this examination, we noted that the AACs have the lowest external drive (not including OLM cells which have zero external drive), and that the NGF cells have either zero or minimal (EWs<1) inputs to them. This can also be easily discerned from the schematic of FIGURE 1 where connection strengths are visualized in a relative fashion. Given this, we considered that PYR cells, PV+BCs, BiCs and CCK+BCs are the critical cell types for the existence of theta rhythms. Focusing on these four cell types, we now move on to develop precise hypotheses of theta-gamma rhythm generation in the FSM. Given the high level of biophysical detail in the FSM, it is reasonable to consider it as a biological proxy. Thus, one can consider extrapolating any determined insights of understanding theta-gamma rhythms and couplings to the biological hippocampal system.

We now zoom in on 300 msec of the full 4 sec raster plot output to do a careful examination of the cell firings in the FSM. This is shown in FIGURE 1C(ii) where only a handful of theta cycles are present and gamma firings can be discerned. We observe certain dynamics that speak to particularly important connections between the four cell types. At the beginning of any given theta cycle the PYR cells start firing as a result of (noisy) external excitation they receive to initiate the theta rhythm. By virtue of their recurrent connectivity, more and more cells can participate and the PYR cell activity (i.e., the number of PYR cells that are active) is elevated due to the progressive recruitment of PYR cells. This activity increase is easily seen in the PYR cell raster plot (FIGURE 1C(ii)). We already know from our previous modeling work that theta rhythm expression is possible with a large enough number PYR cells that have some recurrent connectivity (***Chatzikalymniou et al., 2021***). In that work, we distinguished the PYR cell population, and not any of the inhibitory cell populations, as the theta rhythm initiators. Further, we showed the promotion of increased PYR cell clustering as the recurrent excitation increased (see Figure 6 in ***Chatzikalymniou et al***. (***2021***)). Continuing on, we see that at some critical point the PYR cell activity exceeds a certain threshold that leads to significant activation of the BiCs as well as concurrent enhancement of the PV+BC activity. The strong activation of these PV+ cell types (i.e., PV+BCs and BiCs) in turn strongly inhibits the PYR cells, effectively silencing them. That is, the activity of the PYR cells is sharply terminated once it reaches a threshold due to concurrent PV+ cell activation. Once the PYR cell activity drops, so does the PV+ cell activity which then eventually leads to the PYR cells being able to start firing again. For this, enough PYR cells need to be sitting near their firing threshold. We now evaluate these observations by selectively deactivating connections in the FSM.

### Dissecting the theta/gamma FSM circuit

We test the sharp theta termination directly by removing either PV+BC or BiC connections to the PYR cells. In FIGURE 2A & B, we show that these removals abolish theta rhythms. That is, the theta bursts simply can’t be terminated as the inhibitory connections from PV+BCs or BiCs are no longer there. In FIGURE 2-Supplement 1, we show that theta rhythms are also abolished if the connections from PYR cells to either PV+BCs or BiCs are removed. This is because in the absence of these connections, the PYR cells can’t activate these PV+ cells (BiCs or BCs) once they exceed a critical threshold of activity and as a result the PV+ cells can’t silence them back to sharply terminate their activity and allow another (theta) cycle to begin. With these connection removals, the PYR cells only exhibit a gamma rhythm due to being entrained by the gamma firing inhibitory cells. We note that these simulations in which we removed particular connections expose different aspects relative to the muting experiments done by ***Bezaire et al***. (***2016b***). In Bezaire’s case, muting the cells essentially removed the particular cells from the overall circuit. Here, in removing particular connections, we show the contribution of specific cellular pathways in the generation of theta rhythms.

**Figure 2.**
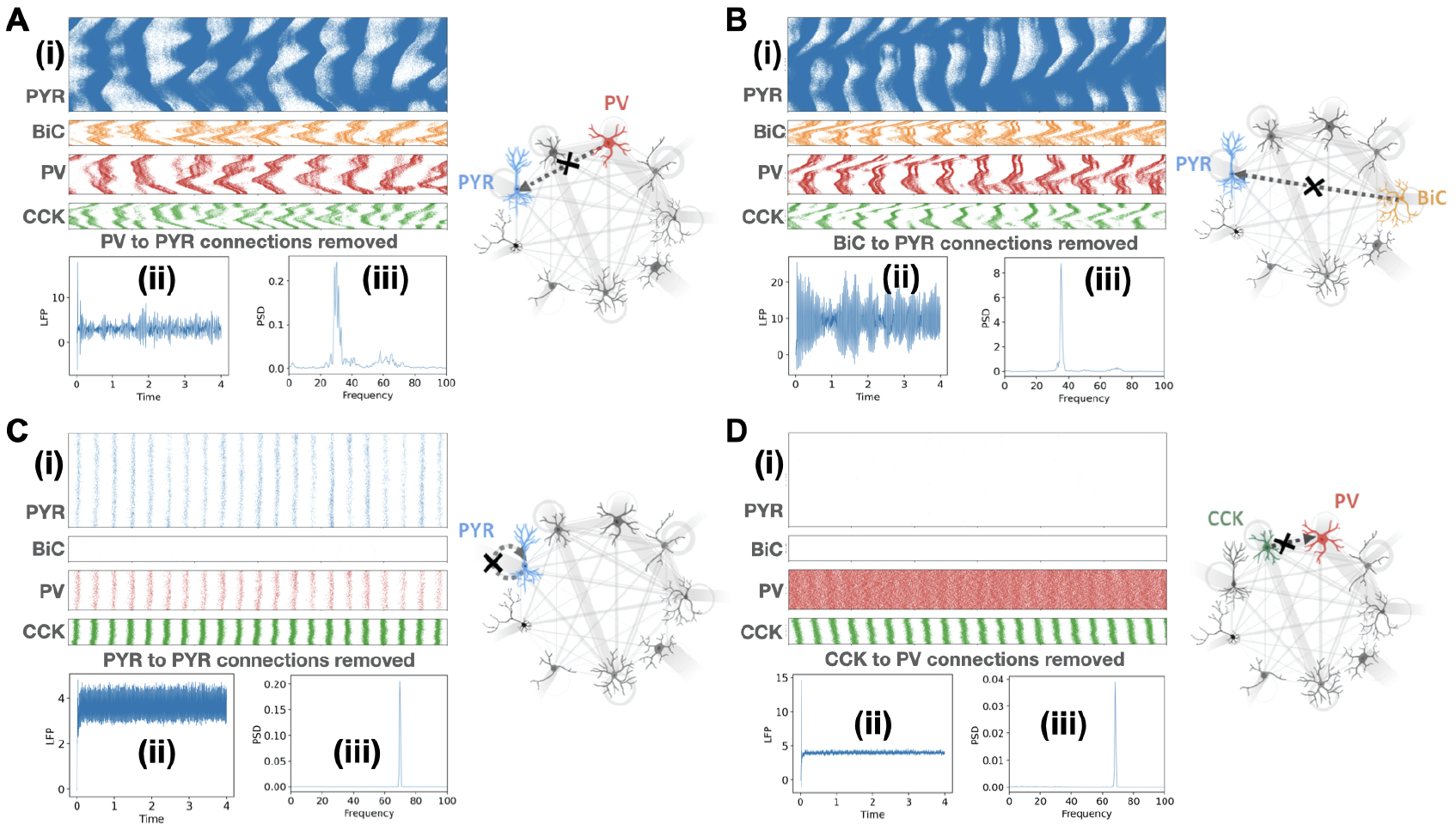
FSM output when connections between different cell types are removed. **A**. PV to PYR connections removed. **B**. BiC to PYR connections removed. **C**. PYR to PYR connections removed. **D**. CCK to PV connections removed. Each part shows a schematic illustration that highlights the particular connections that are removed, and **(i)** show the last 300 ms of raster plot firings for PYR, BiC, PV and CCK cells. **(ii)** unfiltered LFP. **(iii)** PSD of LFP. Units, colors and naming abbreviations are the same as FIGURE 1;

As already noted above, the initiation of the theta rhythm is known to be due to the PYR cell population (***Chatzikalymniou et al., 2021***). The requirement of recurrent connectivity is shown in FIGURE 2C where connections between PYR cells are removed to reveal the loss of theta rhythms. The inhibitory cells contribute to the net input that the PYR cells receive but are not directly responsible for the initiation of the theta rhythm. If there is a reduction in the net input received by the PYR cells and/or the PYR cells are less excitable, then theta rhythms would not be able to be initiated. Interestingly, in our previous modeling we had found that the LFP theta frequency has a linear correlation with the net input received by the PYR cells (***Chatzikalymniou et al., 2021***), and in the FSM of ***Bezaire et al***. (***2016b***), the particular theta frequency is in line with this correlation. Details are provided in the Methods.

Along with PV+BCs and the BiCs, ***Bezaire et al***. (***2016b***) showed that CCK+BCs are pivotal for theta rhythms since there is no theta rhythm with the muting of this cell type. In the FSM there are connections from CCK+BCs to PV+BCs, BiCs and PYR cells. Removing connections from CCK+BCs to BiCs does not eliminate the theta rhythm, but the rhythm is lost if connections from CCK+BCs to either PYR cells or PV+BCs are removed. CCK+BCs provide significant amount of inhibition to the PYR cells and thus removing those connections makes the PYR cells hyper-excitable leading to loss of theta (results not shown). However, the loss of theta with the connection removal to PV+BCs is not so intuitive. We were able to probe this thoroughly by using a ‘clamped’ version of the detailed model network. We did this with a ‘slice’ of the FSM as used before by ***Chatzikalymniou et al***. (***2021***). We performed a grid search of each of the postsynaptic weights received by PV+BCs from the different cell types to find out how these weights impact the PV+BC activity. In particular, we found that a small increase in the synaptic weight from CCK+BCs to PV+BCs could lead to a large decrease in PV+BC firing, akin to a ‘bifurcation point’ in dynamical systems parlance. This was not observed with changes in any of the other synaptic weights from other cell types examined. Details are provided in the Methods. These observations indicate that a strong disinhibition of the PV+BCs, specifically mediated by the CCK+BCs, can induce a sudden increase in the PV+BC firing rate. As shown in FIGURE 2D, this is borne out in the FSM which shows that without CCK+BC to PV+BC connections, there is a large increase in the PV+BC firing rate which in turn shuts down the PYR cells and abolishes theta rhythms. This indicates that CCK+BCs are prominent controllers of PV+BC activity and via that pathway influence theta power.

Looking at the zoomed in raster plots of the focused four cell types in FIGURE 1C(ii), one can clearly discern higher frequency gamma rhythms expressed in PV+BCs and CCK+BC+ as well as in the PYR cells where the slower frequency theta rhythms are also apparent. Theta and gamma rhythms are both present in the LFP signal as seen from the power spectrum (FIGURE 1B(ii)). The gamma rhythm seen in the LFP output is produced by the gamma-paced entrainment of the noisyfiring PYR cells as follows: The PV+BCs and the CCK+BCs fire at gamma frequencies and organize themselves into coherently firing populations. The PV+BCs and the CCK+BCs also form connections with the PYR cells. As a result, during a theta cycle these gamma-firing cells periodically inhibit the noisy firing PYR cells forcing them to burst in a gamma-paced manner. This results in a gamma rhythm being present with the progressive recruitment of PYR cells toward the peak of the theta cycle. Because the amplitude of the gamma LFP maximizes at peak of the PYR cell theta burst (which corresponds to the peak of the theta cycle), the amplitude of the gamma rhythm is coupled with the phase of theta, giving rise to phase-amplitude coupling of the theta and gamma rhythms.

### A reduced circuit model

From the FSM simulations and analyses described above we now have precise cell-based hypotheses of how theta/gamma rhythms are generated in the CA1 hippocampus. Namely, the theta rhythm is initiated by the PYR cell population, sharply terminated by PV+BCs and BiCs, and connections from CCK+BCs to PV+BCs are essential for theta rhythms. By removing particular connections in the FSM, these hypotheses were tested and confirmed. Given this, we feel confident to consider a reduced circuit model that is derivative of the FSM. This reduced model consists of the four different cell types and only those connections deemed essential in light of the developed hypotheses. A schematic of this reduced circuit model relative to the FSM is shown in FIGURE 3. This reduced circuit model motivates the creation of a population rate model (PRM).

**Figure 3.**
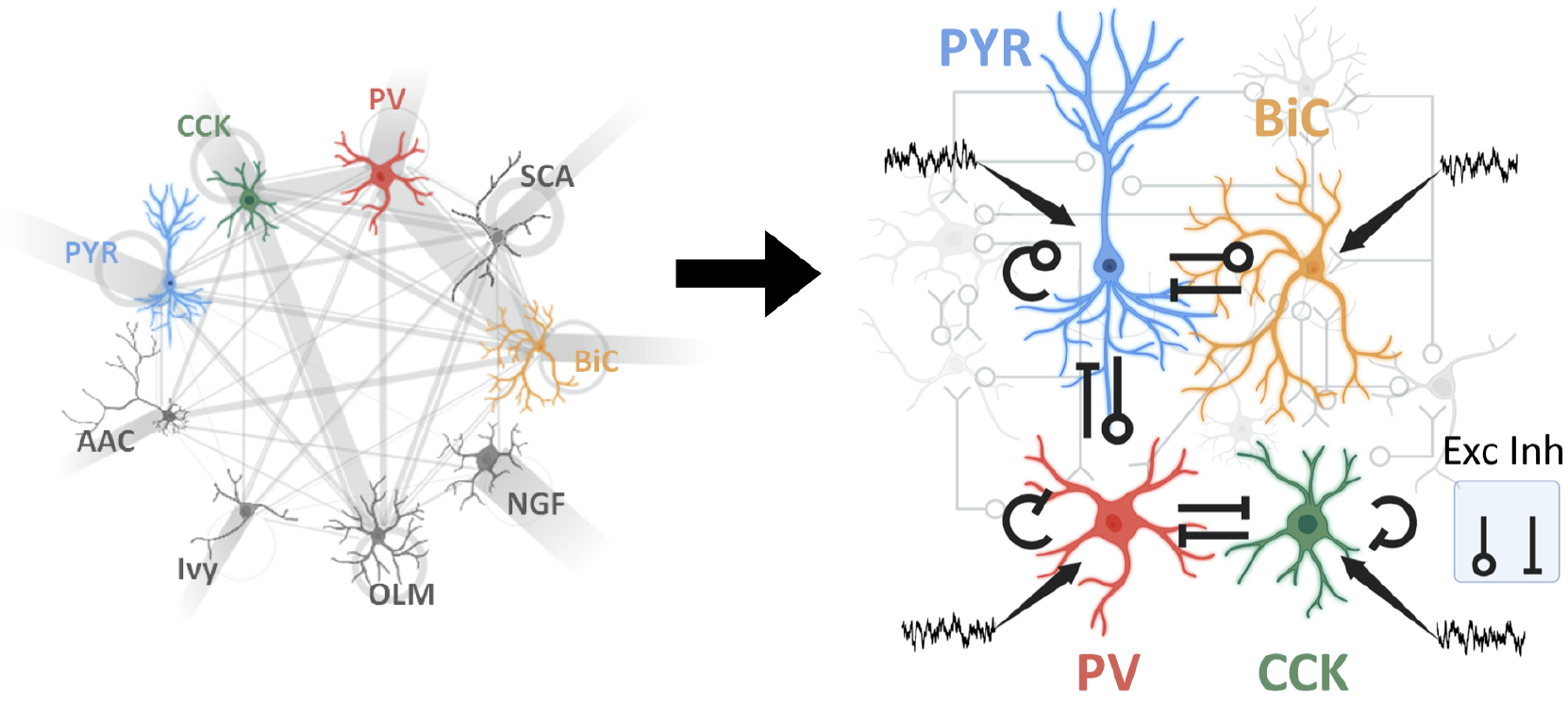
Rationalized reduction in complexity of CA1 hippocampus model circuit. The left side shows the stylized schematic of the FSM. Detailed descriptions of color and naming abbreviations are given in FIGURE 1. The right side shows a stylized schematic of the reduced model with 4 cell types and excitatory (Exc) and inhibitory (Inh) connections between them and inputs (incoming arrows). The rationale used to consider this reduced model is given in the main text.

### Population rate model (PRM) reproduces FSM derived hypotheses

Considering the reduced circuit model shown in FIGURE 3, we set up a population rate model (PRM) that is able to exhibit multi-modal (theta/gamma) oscillations. The equations are provided in the Methods. Using this PRM we found a set of parameter values that give rise to theta-gamma rhythms as shown in FIGURE 4A. We refer to these parameter values as ‘reference parameters’, and the values are provided in the Methods (TABLE 4). These theta-gamma rhythms are in accordance with the developed hypotheses. That is, the presence of theta rhythms requires PV+BC and BiC to PYR cell connections as well as CCK+BC to PV+BC connections. In line with the hypotheses, theta rhythms are lost when these various connections are removed, as in FIGURE 4B-D. It is interesting that this reference parameter set produces activities of CCK+BCs and PV+BCs that fire alternately as has been experimentally shown (***Dudok et al., 2021a***). We now undertake a full-blown parameter exploration of the PRM and perform analyses and visualizations of thousands of simulations to obtain the following insights and predictions.

**Figure 4.**
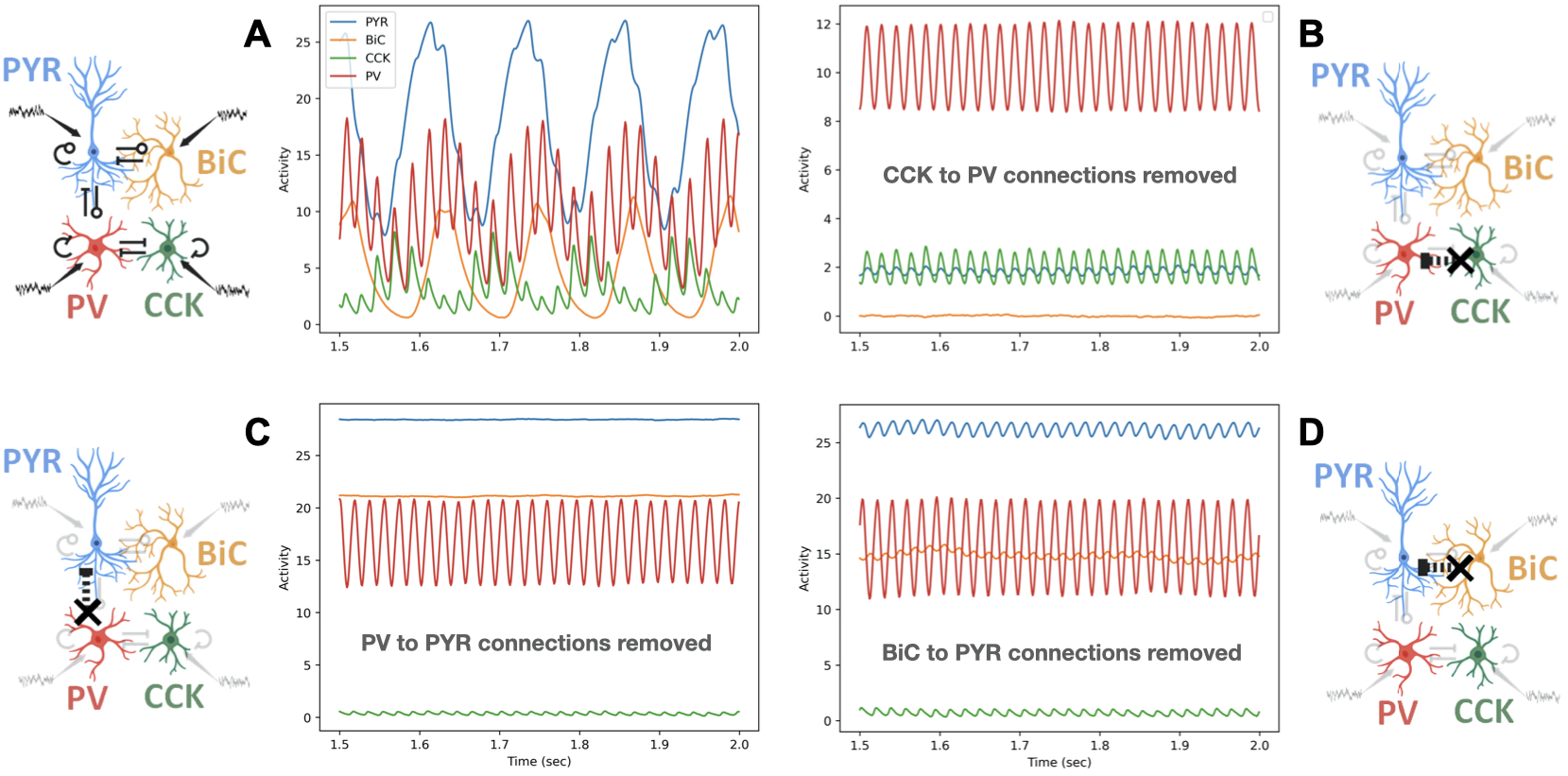
Output of the reduced population rate model (PRM). **A**. PRM output of the activity of each cell type population (see TABLE 4 for parameter values). **B**. PRM output when CCK to PV connections are removed. **C**. PRM output when PV to PYR connections are removed. **D**. PRM output when BiC to PYR connections are removed. Naming and color scheme of the 4 cell types is the same as in FIGURE 1. Each part shows a schematic illustration that highlights the particular connections that are removed.

**Table 4.** Reference parameter values for the PRM.

| Symbol | Definition | Value |
| --- | --- | --- |
| $\beta$ | Response function gain | 10au |
| $\tau$ | Conduction delay | 5ms |
| $\alpha_{PYR}$ | PYR cell rate constant | 50Hz |
| $\alpha_{BiC}$ | BiC rate constant | 50Hz |
| $\alpha_{CCK}$ | CCK+BC rate constant | 80Hz |
| $\alpha_{PV}$ | PV+BC rate constant | 100Hz |
| $w_{PYR \rightarrow PYR}$ | $PYR \rightarrow PYR$ coupling | 0.03au |
| $w_{PYR \rightarrow BiC}$ | $PYR \rightarrow BiC$ coupling | 0.04au |
| $w_{PYR \rightarrow PV}$ | $PYR \rightarrow PV$ coupling | 0.02au |
| $w_{BiC \rightarrow PYR}$ | $BiC \rightarrow PYR$ coupling | -0.03au |
| $w_{CCK \rightarrow CCK}$ | $CCK \rightarrow CCK$ coupling | -0.15au |
| $w_{CCK \rightarrow PV}$ | $CCK \rightarrow PV$ coupling | -0.15au |
| $w_{PV \rightarrow PV}$ | $PV \rightarrow PV$ coupling | -0.055au |
| $w_{PV \rightarrow PYR}$ | $PV \rightarrow PYR$ coupling | -0.04au |
| $w_{PV \rightarrow CCK}$ | $PV \rightarrow CCK$ coupling | -0.075au |
| $i_{PYR}$ | Input to PYR cells | 0.07au |
| $i_{BiC}$ | Input to BiCs | -1.05au |
| $i_{CCK}$ | Input to CCK+BCs | 0.7au |
| $i_{PV}$ | Input to PV+BCs | 0.45au |
| $D_{PYR}$ | Noise variance to PYR cells | 0.001au |
| $D_{BiC}$ | Noise variance to BiCs | 0.001au |
| $D_{CCK}$ | Noise variance to CCK+BCs | 0.001au |
| $D_{PV}$ | Noise variance to PV+BCs | 0.001au |
| $r_o$ | Maximum firing rate | 30Hz |

### PRM predictions are in agreement with FSM predictions on theta frequency control and the requirement of recurrent collaterals

In our previous work we showed that the LFP population theta frequency is correlated with the net input received by the PYR cells (***Chatzikalymniou et al., 2021***). Further, this correlation was found to be mostly due to the excitatory external drive and the inhibitory input, and not mainly due to recurrent collateral inputs - see Figure 7d(i-iii) in ***Chatzikalymniou et al***. (***2021***). We found that as excitatory or inhibitory input to PYR cells increased, the theta frequency increased. The 2-D heat maps of FIGURE 5 show that this is also the case with the PRM. Specifically, increasing the excitatory input to PYR cells (*i*_*PYR*_) led to increases in the theta frequency as shown in FIGURE 5A(i) where *i*_*PYR*_ varies along the x-axis. Increasing the inhibitory input in the form of larger connection weights (i.e., more negative values) from BiCs (*w*_*BiC*→*PYR*_) or from PV+BCs (*w*_*PV*→*PYR*_) to the PYR cells also resulted in theta frequency increases. This is shown in FIGURE 5B(i) & C(i) where *w*_*PV*→*PYR*_ and *w*_*BiC*→*PYR*_, respectively, vary along the x-axis of the 2-D heat maps. However, modulating the recurrent excitation (*w*_*PYR*→*PYR*_) did not produce changes in theta frequencies. To show this more directly, we plot theta frequency versus the different parameters in 1-D plots in FIGURE 5A(ii). This plot shows that there is no change in the theta frequency as the recurrent excitatory weight (*w*_*PYR*→*PYR*_) is changed (blue triangles), but that there is a frequency increase as the excitatory drive *i*_*PYR*_ is increased (red circles). Additional plots in FIGURE 5B(ii) & C(ii) show that as inhibitory input to the PYR cells increases, due to either increasing synaptic weights from BiCs (*w*_*BiC*→*PYR*_) (blue circles in C(ii)) or from PV+BCs (*w*_*PV*→*PYR*_) (green circles in B(ii)), the theta frequency increases. The particular values shown in these plots are shown as horizontal or vertical lines (red part of the lines) in the 2-D heat maps of FIGURE 5A(i), B(i) & C(i).

**Figure 5.**
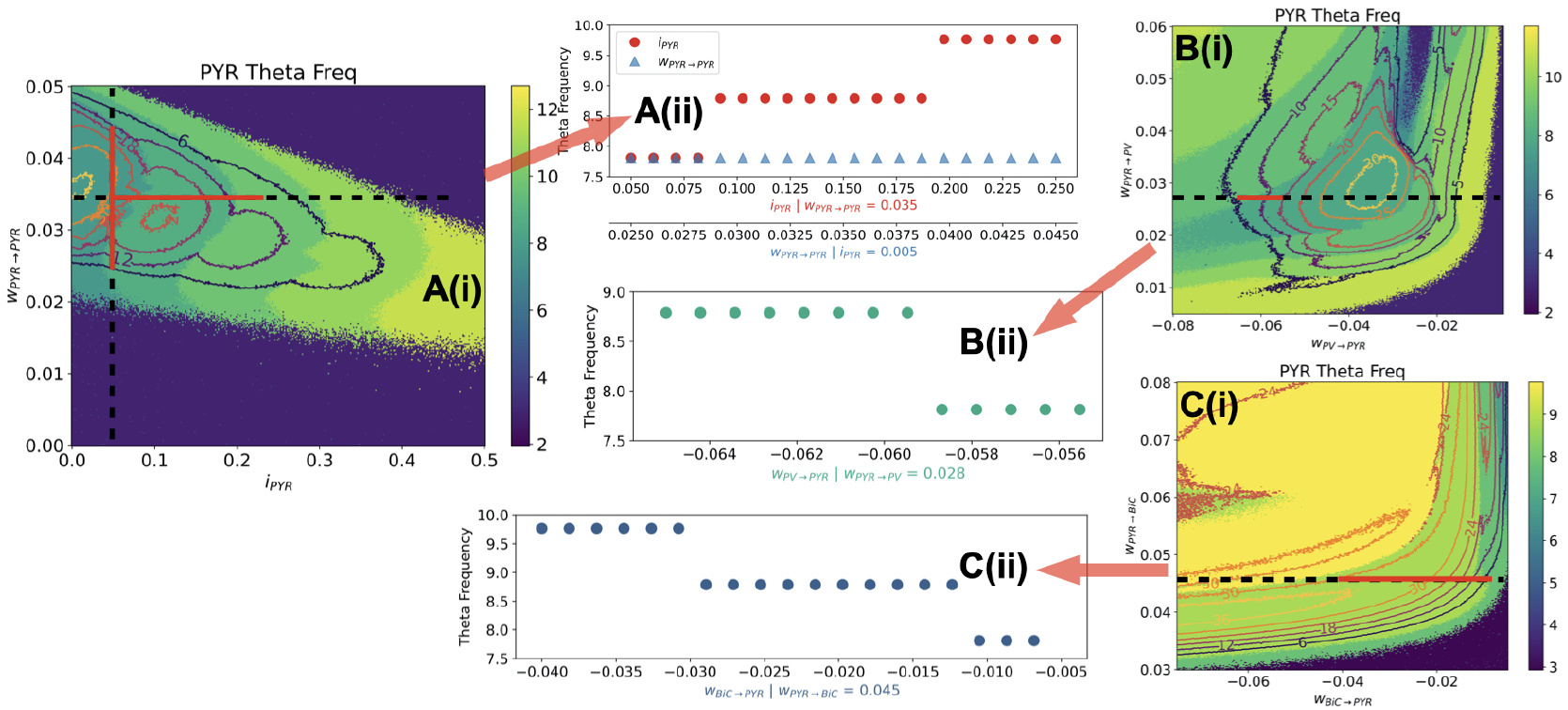
Theta Frequency Control in PRM. Theta frequency changes in the PYR cells for changes in parameter set pairs. **A**. Heat map for (*i*_*PYR*_ − *w*_*PYR*→*PYR*_) is shown in **(i)** and a plot from a slice is shown in **(ii). B**. Heat map for (*w*_*PV*→*PYR*_ − *w*_*PYR*→*PV*_) is shown in **(i)** and a plot from a slice is shown in **(ii). C** Heat map for (*w*_*BiC*→*PYR*_ − *w*_*PYR*→*BiC*_) is shown in **(i)** and a plot from a slice is shown in **(ii)**. The contours in the heat maps encompass the 6 highest power values, i.e., where there are high activity values at theta frequencies. ‘Slices’ of the heat maps are from the indicated dashed black lines, and the plots are specifically shown for ranges indicated by the red part of the dashed black lines, as chosen to more easily see the trends in theta frequency changes.

Given the different and reduced structure of the PRM, we do not expect a direct correspondence between PRM and FSM-generated frequencies. However, it is reassuring that we obtain similar relationships - that is, increases or decreases in theta frequency as previously predicted using the detailed model (***Chatzikalymniou et al., 2021***). This implies that usage of this PRM can produce valid relationships relative to the biological system as represented by the detailed model. We do note that there is much richness in the PRM that can be exposed due to the expansive exploration made possible by its much reduced nature relative to the detailed model. For example, there are parameter regimes in which the theta frequency decreases (rather than increases) when *w*_*PV*→*PYR*_ connection weights are increased - specifically, for the lower weight (less negative) values of *w*_*PV*→*PYR*_. This can be seen in the particular heat map of FIGURE 5B(i). In FIGURE 5-Supplement 1, we show the heat maps of theta frequency for the other three cell types (PV+BCs, BiCs, CCK-BCs).

As was previously shown (***Bezaire et al., 2016b***; ***Chatzikalymniou et al., 2021***; ***Ferguson et al., 2017***), recurrent connections (*w*_*PYR*→*PYR*_) are needed for the existence of theta rhythms. To visualize this, we compute 2-D difference heat maps that represent the subtraction of the normalized gamma power from the normalized theta power. Blue is used to show where theta rhythms dominate, and red where gamma rhythms do. See Methods for details regarding these computations. In this way, we are able to nicely visualize how different parameters affect theta and gamma rhythms and their coupling. In FIGURE 5-Supplement 2, we show difference heat maps for a wide range of *i*_*PYR*_ − *w*_*PYR*→*PYR*_ parameter values, and from it, it is abundantly clear that a non-zero value of *w*_*PYR*→*PYR*_ is needed to have theta rhythms. In FIGURE 5-Supplement 2, we also show a plot of the activities of the four different cell types when there is no recurrent excitation - theta rhythms are not present.

### PRM predicts stronger control by CCK+BCs relative to PV+BCs for theta vs gamma rhythm dominance

In FIGURE 6A & B, we respectively show 2-D difference heat maps of *i*_*CCK*_ − *i*_*PV*_ and *w*_*PV*→*CCK*_ − *w*_*CCK*→*PV*_ parameter sets. FIGURE 6A(i) & B(i) maps show parameter ranges that were chosen to clearly show regimes of theta (blue) or gamma (red) rhythm dominance, whereas FIGURE 6A(ii) & B(ii) maps show parameter ranges that are the same for both x- and y-axes. Between the blue and red regions is a yellow region where both theta and gamma rhythms are present - i.e., theta-gamma coupling exists. This theta-gamma coupling is easily seen in the activities of the four cell types in FIGURE 4A for the reference parameters. In particular, in the PYR cell (blue) activity, but also in the PV+BC (red) and CCK+BC (green) activities. To more easily visualize this coupling regime, we add a black line to the difference heat maps of FIGURE 6A & B where switching between gamma or theta dominance would occur, that is, where there is theta-gamma coupling. What is abundantly clear from the steepness of the black lines in these difference heat maps is that CCK+BCs relative to PV+BCs can more strongly control the level of theta versus gamma rhythm dominance. For example, a small change in the amount of input being received by CCK+BCs from PV+BCs (*w*_*PV*→*CCK*_ in FIGURE 6B(i),B(ii)) can switch the dominance of whether theta or gamma rhythms are expressed. This would not be the case for a similar small change in the amount of input being received by PV+BCs from CCK+BCs (*w*_*CCK*→*PV*_). This stronger control of theta-gamma coupling by CCK+BCs relative to PV+BCs is less obvious in the difference heat maps when the drive to either CCK+BCs (*i*_*CCK*_) or PV+BCs (*i*_*PV*_) (FIGURE 6A(ii)) is considered for the same parameter values. That is, the slope of the black line on the *i*_*CCK*_ versus *i*_*PV*_ difference heat map (FIGURE 6A(ii)) is less steep than that of the black line on the *w*_*PV*→*CCK*_ versus *w*_*CCK*→*PV*_ difference heat map (FIGURE 6B(ii)). In FIGURE 6-Supplement 1, we show the difference maps of theta frequency from the perspective of the other three cell types (PV+BCs, BiCs, CCK-BCs). Difference heat maps with PYR and PV+BC couplings further support this as additional ‘input’ received by PV+BCs from PYR cells does not sensitively (i.e., steeply) affect theta-gamma coupling - see FIGURE 6-Supplement 2. Overall, it is clear that the CCK+BCs more strongly control theta-gamma coupling relative to the PV+BCs as observed from the different effects of input changes to CCK+BCs or PV+BCs.

**Figure 6.**
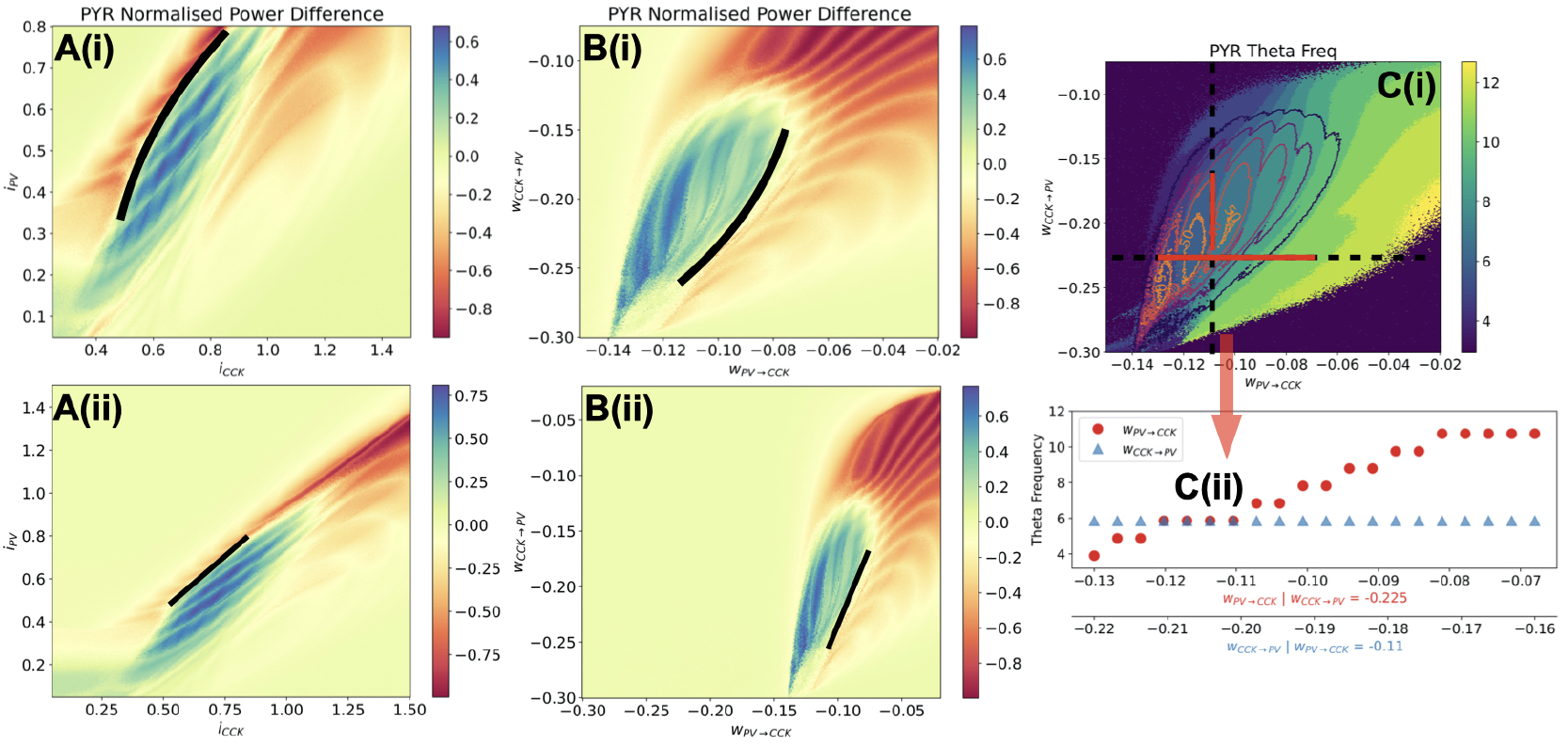
CCK+BCs more strongly control rhythm expression and theta frequencies. Difference Maps: [(Normalized power of theta) - (Normalized power of gamma)] are shown in **A** and **B**. Positive (blue) values mean that theta rhythms are dominant, and negative (red) values mean that gamma rhythms are dominant. ‘Yellow’ values mean that theta and gamma powers are comparable so that there is theta-gamma coupling (or that both theta and gamma powers are very small). The difference map plots show CCK+BCs and PV+BCs from drive **A(i)** and connection perspectives **B(i)**, and when viewed for the same parameter range values **A(ii)** and **B(ii)**. Black lines are added to delineate theta and gamma dominant regions. Given the angle of these lines, it is clear that CCK+BCs have a higher level of control relative to PV+BCs in determining whether theta or gamma rhythms are dominant, and hence coupled, since the slopes are steeper for changes from CCK+BC perspectives. **C(i)** shows the theta frequency heat map for *w*_*PV*→*CCK*_ − *w*_*CCK*→*PV*_ connection weights and a plot of the theta frequency to show that PV+BC to CCK+BC connection more strongly affect the theta frequency is shown in **C(ii)**. As in Figure 5, the dashed black lines show the ‘slices’ taken from the heat map in plotting the theta frequency versus either *w*_*PV*→*CCK*_ (red dots) or *w*_*CCK*→*PV*_ (blue trianges) for ranges indicated by the red part of the dashed black lines.

An interesting observation is that the theta frequency is more strongly affected by modulating connection strengths from PV+BCs to CCK+BCs than the other way. This is visualized in the frequency heat map shown in FIGURE 6C(i). Similar to FIGURE 5, we show a theta frequency plot (FIGURE 6C(ii))) as these connections are varied, with horizontal or vertical dashed line representing the particular values in the plot where *w*_*PV*→*CCK*_ (horizontal) or *w*_*CCK*→*PV*_ (vertical) changes occur. It is clear that the theta frequency minimally changes with *w*_*CCK*→*PV*_ modulation, unlike with *w*_*PV*→*CCK*_ modulation. In FIGURE 6-Supplement 1 SUPP FIG 4, we show the heat maps of theta frequency for the other three cell types (PV+BCs, BiCs, CCK-BCs).

The prediction of the importance of CCK+BCs for theta-gamma coupling from the PRM was possible because of the extensive and systematic parameter exploration that could be carried out in the reduced circuit model (FIGURE 3). We note that this importance of CCK+BCs in population rhythms is in line with observations from our detailed model explorations described above, but from a different perspective. We had showed that CCK+BCs, unlike any of the other cell types, could drastically affect PV+BC firing which in turn strongly suggests that they underlie the control of theta expression. Indeed, we showed that theta rhythms are not present when CCK+BC to PV+BC connections are removed in the FSM (see FIGURE 2D). Experimentally, CCK+BCs and PV+BCs show complementary activities and CCK+BCs inversely scale with PYR cell activities ***Dudok et al***. (***2021a***) This would be in line with disinhibition of PV+BCs by CCK+BCs and its subsequent effect on PYR cell firing. Additionally, ***Fasano et al***. (***2017***) have shown that CCK+BCs, as identified by atypical vesicular glutamate transporters, alter theta oscillations.

## Discussion

In this paper, we presented a detailed explanation of how theta and gamma rhythm mechanisms could arise in the CA1 hippocampus and showed that CCK+BCs play a controlling role in theta-gamma coupled rhythms. We built upon earlier modeling studies that brought together minimal (***Ferguson et al., 2017***) and detailed (***Bezaire et al., 2016b***) CA1 microcircuit models with defined inhibitory cell types that produce theta rhythms and showed that large enough PYR cell networks could initiate theta rhythms with inhibitory cell populations ‘regularizing’ the rhythm. We termed this an ‘inhibition-based tuning’ mechanism (***Chatzikalymniou et al., 2021***; ***Skinner et al., 2021***). The theta rhythm generation is initiated by the PYR cell population which also sets the theta frequency, but the termination of the theta bursts is an inhibition-mediated mechanism which contributes to the robustness and stability of the rhythm. In their original study, ***Bezaire et al***. (***2016b***) described the preferential discharge of the PYR cells at the trough of the LFP analog and the subsequent strong recruitment of PV+BCs and BiCs to silence the PYR cells, but the specifics leading to the termination of the theta burst were not parsed out. They also described the progressive recruitment of PYR cell firings towards the peak of the theta cycle, but the prominent role of the PYR cells as theta rhythms initiators was first demonstrated and discussed in ***Chatzikalymniou et al***. (***2021***).

From our explorations of the detailed model here, we directly showed that PV+BCs and BiCs are responsible for the sharp termination of PYR cell firing. Of note, we found that CCK+BCs, more than any other inhibitory cell type, play a key role in PV+BC firing. Gamma rhythms arose due to coherent firing in CCK+ and PV+ basket cell networks. Using the four cell types of PYR cells, PV+BCs, CCK+BCs and BiCs, we generated theta-gamma rhythms in a reduced population rate model (PRM). The PRM reproduced the specific contributions of these four different cell populations seen in the detailed full-scale model (FSM). From extensive simulations and visualization analyses of the PRM, we predicted that CCK+BCs more strongly controlled theta-gamma coupled rhythms than PV+BCs. We further predicted that PV+BC to CCK+BC connections can more strongly affect theta frequencies relative to CCK+BC to PV+BC connections. While we were able to capture much with our many 2D visualization plots from our thousands of simulations, further insights may be possible by doing theoretical analyses of the PRM. This type of extensive exploration can be done using such reduced models that have much less parameters relative to detailed models. Importantly, as the PRM was built based on precise hypotheses developed from the biophysically detailed FSM, the predictions arising have more biological plausibility regarding cell type specifics, relative to a ‘generic’ PRM. As tools to identify and target CCK+BCs have been developed (***Dudok et al., 2021a***), we can envisage designing experiments to directly test predictions arising from our work.

The initiation of theta rhythms by the PYR cell population in the hippocampus is also likely to be the case in the biological system since the existence of ‘threshold behaviour’ in initiating hippocampus population bursts has been shown before by ***de la Prida et al***. (***2006***). Also, theta-resonant behaviour is expressed by PYR cells (***Hu et al., 2002, 2009***). Interestingly, in another network modelling study, the necessity of PYR cells to be near their threshold for firing was exploited to explain how reduced excitatory fluctuations could lead to hyper-excitability in mouse models of Rett syndrome (***Ho et al., 2014***).

The high frequency gamma rhythms seen in the model LFP (entwined with the slower frequency theta rhythm) is more akin to the so-called ING (interneuron network gamma) rather than PING (pyramidal-interneuron network gamma) mechanism (***Whittington et al., 2000***). That is, the PV+ and CCK+ BC networks have appropriate ‘input and balance’ to create coherent network output at gamma frequencies, which is then imposed onto the PYR cell population. It is not a PING mechanism since gamma coherence does not (overall) rely on phasic input from PYR cells. The fact that there is a phasic aspect to the theta/gamma rhythms in the model is related to the described PYR cell clustering aspects. PING and ING certainly help us think about what sort of excitatory and inhibitory balances might be key, but the additional cell types and the multi-modal oscillation considerations here supercede these classical ideas. Naturally, what ultimately matters is what experiments can be done to differentiate and understand the contributions from the various excitatory and inhibitory cell types in producing the circuit output, such as considered for visual cortex (***Tiesinga and Sejnowski, 2009***).

Our results are of course limited by what exists in the detailed FSM which was put together as carefully as possible given a ‘knowledge base’ of the available experimental data at the time (***Bezaire and Soltesz, 2013***). For example, we did not try to include PYR cell to CCK+BC connections in the PRM since they are not present in the FSM even though such connections possibly exist (***Dudok et al., 2021a***). Also, we did not include CCK+BC to PYR cell connections in the PRM even though they are present in the FSM. This does not negate the PRM results here. Rather, our interpretation would be that CCK+BC to PV+BC connections are more critical rhythm controllers than connections from CCK+BCs to PYR cells. We note that the AACs form strong connections with the PYR cells in the FSM and it is likely that they contribute to the termination of the theta burst along with the PV+BCs and BiCs by providing adequate inhibition and contributing to the excitatory-inhibitory balance necessary for theta rhythms (***Chatzikalymniou et al., 2021***).

While the theta mechanism presented in this study is based on the specific connectivity of the FSM network, the relative strength of connections between inhibitory populations evidently changes across the septo-temporal axis of the CA1 and across layers (***Soltesz and Losonczy, 2018***; ***Navas-Olive et al., 2020***). ***Navas-Olive et al***. (***2020***) have shown that PV+BCs preferentially innervate PYR cells at the deep sublayers while CCK+BCs are more likely to target the superficial PYR cells. Also, there is different innervation of PYR cells by BiCs and OLM cells in deep versus superficial PYR cells. How these differential connectivities affect theta-gamma rhythm control could be understood by examining various dynamic balances in the PRM. Further, specific known motifs that include VIP+ cell types (***Guet-McCreight et al., 2020***) are indirectly encompassed in the PRM via the inputs received by the different cell types and so their contributions can be indirectly considered in the theta/gamma mechanisms. Moving forward, one could imagine expanding the PRM to include additional cell type populations and connectivities to develop further hypotheses and to obtain predictions for experimental examination.

The plethora of inhibitory cell types with particular biophysical characteristics and connectivities in the hippocampus (***Pelkey et al., 2017***) make it challenging to figure out their particular contributions to dynamic network oscillations in the behaving animal. As theta and gamma oscillations and their coupling reflect cognitive processing and are potential disease biomarkers, *and* involve the various inhibitory cell types, we cannot ignore the potential roles played by these different types of interneurons. Interestingly, it has been shown that timescale gradients are significantly correlated with inhibitory cell-type markers (***Gao et al., 2020***). In conclusion, our work presented here shows that we can determine specific contributing roles of various inhibitory cell types by developing hypothesis-driven population rate models from detailed biophysical models. As tools to identity and control particular inhibitory cell types become more available, one can imagine leveraging these tractable PRM models for prediction and experimental design.

## Methods

### Detailed CA1 microcircuit: full-scale model (FSM)

The original fullscale CA1 microcircuit repository which can be found at ModelDB at: https://senselab.med.yale.edu/ModelDB/showModel.cshtml?model=187604. A new model version in python is now also available at: https://github.com/dhadjia1/ca1-microcircuit. Analysis of simulation outputs can be recreated using the publicly available SimTracker tool (***Bezaire et al., 2016a***). Documentation on SimTracker and installation instructions can be found at: https://simtrackercode.readthedocs.io/en/compiled/. It is recommended that users install SimTracker first and then install and register the CA1 model under SimTracker, to take advantage of the visualization functionalities of the SimTracker package. This tool is offered both as a stand-alone, compiled version for those without access to MATLAB (for Windows, Mac OS X, and Linux operating systems), and as a collection of MATLAB scripts for those with MATLAB access. Once the SimTracker and the CA1 repository are installed, users can run simulations either on their local machines using a small scale of the CA1 network, or on supercomputers as needed for full scale network simulations. To reproduce the findings presented here, one needs to first familiarize oneself with the CA1 microcircuit background and code.

A 10% ‘slice’ of the FSM (***Bezaire et al., 2016b***) is used in ***Chatzikalymniou et al***. (***2021***). It is created from the FSM by reducing the volume and the number of cells by a tenth, as well as dividing all the connections in the network by a factor of ten so that we have a ‘slice’ and not a normalization of the FSM. The grid search is carried out using this model. In the FSM there are eight different inhibitory cell types and excitatory PYR cells. All of these cell types are connected in empirically specific ways based on an extensive knowledge-based review of the literature (***Bezaire and Soltesz, 2013***). The cells are evenly distributed within the various layers of the CA1 (stratum lacunosum-moleculare, radiatum, pyramidale, oriens) in a three-dimensional prism. Afferent inputs from CA3 and entorhinal cortex (EC) are also included in the form of Poisson-distributed spiking units from artificial CA3 and EC cells (referred to as ‘other input’ in TABLE 1). We note that although there are layer-dependent specifics regarding how the different cell types are arranged in the FSM, there are not any differences along the longitudinal axis. As such, the connection probabilities in any particular part of the longitudinal axis would be the same. Power spectral densities of FSM LFP output shown in FIGURES 1-2 are generated using a Welch periodogram python code -the same as that used in analyzing the population rate model described below.

#### Input currents to individual cells of the FSM

In order to obtain the input currents received by a cell in the FSM, the Network Clamp tool of the SimTracker software package (***Bezaire et al., 2016a***) was used. Using data obtained from the full-scale CA1 microcircuit simulations of ***Bezaire et al***. (***2016b***) that are available on the CRCNS repository at: https://portal.nersc.gov/project/crcns/download/index.php, and the Network Clamp tool, one can explore the activity of any single neuron within the network. In this manner, we investigated five PYR cells and the input currents each of them received from all the other cell types in the network in the control network. We computed the mean net input current (−9.35 nA) across these five PYR cells, as well as the standard deviation in the net input current (0.13 nA). From a periodogram of the FSM of Bezaire et al, the theta frequency is 8.5 Hz. From the linear correlation between theta frequency and input current that we previously obtained (***Chatzikalymniou et al., 2021***), this frequency value would be due to an input current around 10 nA. Thus, the computed net input current here reasonably agrees with this relationship.

### Grid search details using the network clamp technique

Using the Network Clamp tool, we can take a snapshot of the incoming synaptic input received by any particular cell in the network. We used it to undertake an in-depth exploration of the control of PV+BC activity due to inputs from other cell types. We applied it as follows: First, we obtained the output of an initial full-network simulation and ran network clamp simulations on a PV+BC. We altered the incoming afferent synapse weights (but not the incoming spike trains) and examined how PV+BC activity was affected. This approach gives an estimate of how the synaptic weights affect the PV+BC firing. From this, one can gain insight into how the post-synaptic weights might affect the full network simulations.

The manner in which we undertook our exploration is illustrated in FIGURE 7. We used the following pipeline of analysis:

**Figure 7.**
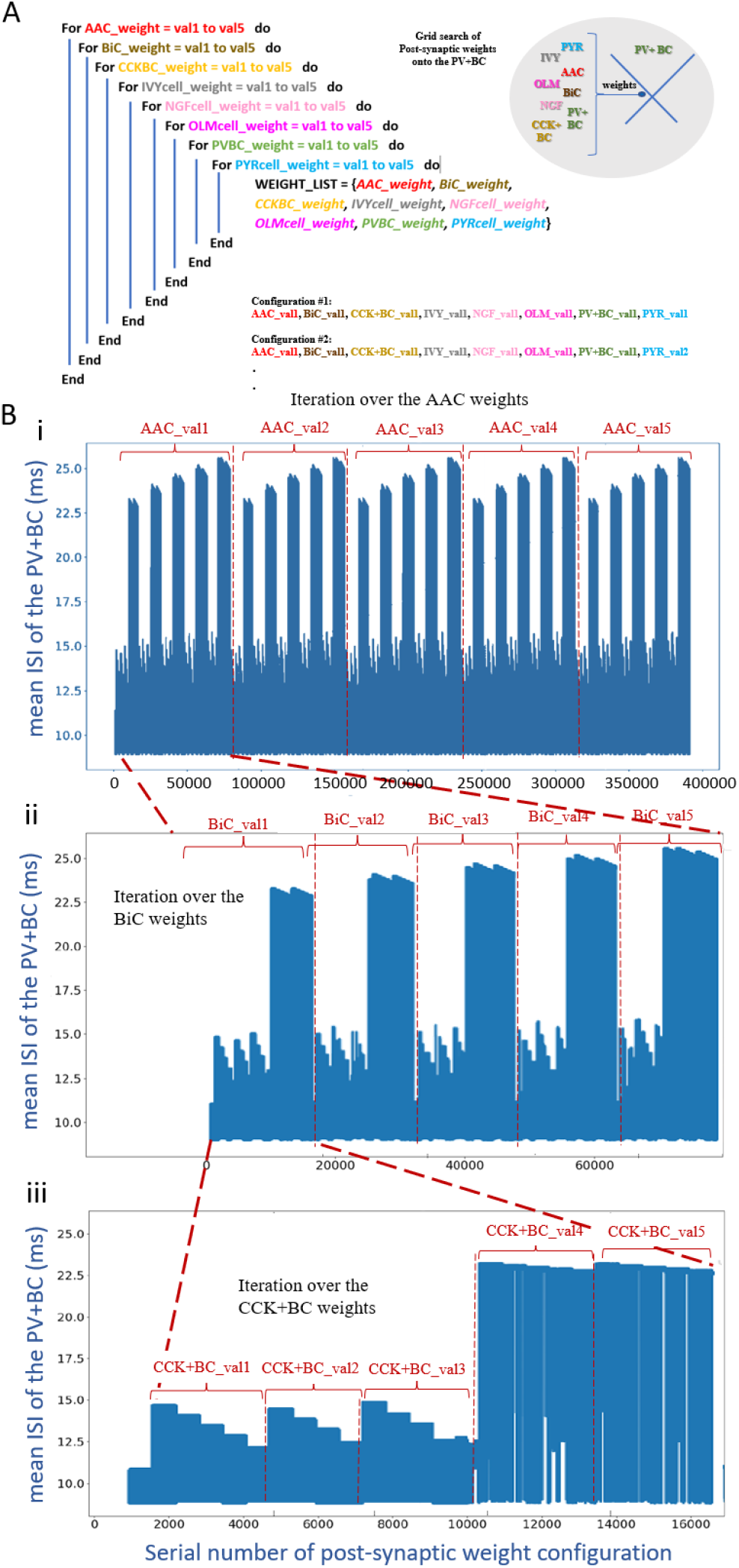
The PV+BC ISIs as a function of the configuration number of the post-synaptic weight combinations. We iterate over the post-synaptic PV+BC weights according to the process described Methods. **A**. Recursive algorithm of the grid search over a range of five post-synaptic weights per cell type. **B**. Iteration over the postsynaptic weights gives rise to a fractal-like pattern in which similar patterns recur at progressively smaller scales. (i) On the largest scale we identify a group of five repetitive motifs where each motif corresponds to a given AAC post-synaptic weight value. (ii) Within each of these motifs another group of five repetitive motifs corresponds to the five BiC post-synaptic weight values. (iii) Equally within each one of these five motifs another group of five similar motifs corresponds to the five CCK+BC post-synaptic weight values and so on. Incremental increases of the other five post-synaptic cell weights which don’t influence PV+BC firing significantly.

1. Explore the post-synaptic weights onto a given PV+BC. Starting from a given set of reference values (r.v.), vary the post-synaptic weights from all eight pre-synaptic cell types to the PV+BC, and search the range of post-synaptic weights for each pre-synaptic cell type. The range used is [-50% (r.v.), +50% (r.v.), 25%] (*start value, end value, step*), and as such it contains 5 values which are: −50% (r.v.) | −25% (r.v.) | r.v. | +25% (r.v.) | +50% (r.v.). Explore the co-variation of these values. Since every post-synaptic weight can take 5 values and that we explore a set of eight post-synaptic weights, we have a total of 5^8^ = 390,594 configurations.
2. Perform network clamp simulations for all configurations and store the voltage trace of the PV+BC for each configuration.
3. Calculate the inter-spike intervals (ISIs) of every voltage trace using the open-source eFEL library (https://github.com/BlueBrain/eFEL).
4. Plot the mean ISI across all configurations. This is shown in FIGURE 7.

## Observations

In FIGURE 7, we show the result of a grid search representing the co-variation of eight post-synaptic conductances. Following the simple visualization scheme shown in FIGURE 7A, we can easily discern the effect that each presynaptic cell has on the firing rate of the PV+BC. We distinguish a number of patterns. As expected, incrementally strengthening the inhibitory PV+BCs post-synaptic weights decreases the PV+BC firing rate (larger ISIs). However, for a critical value of the CCK+BC to PV+BC conductance, we noticed a sudden decrease in the PV+BCs firing frequency which can be up to 20Hz. This transition always occurred at a critical value of the CCK+BC to the PV+BC conductance. Once that critical value is reached, a small increase in the rest of the conductances, especially the PV+BC to PV+BC, mediates an abrupt elevation of the PV+BC mean firing rate (by means of ISI). This observation reveals that, provided that the network operates close to this critical point, small changes in the CCK+BC activity can induce a sudden change (increase or decrease) of the PV+BC firing rate. Such a critical value always exists regardless of the combination of the rest of the post-synaptic conductances.

In essence, our grid search shows that if one sits close to a critical point (a point representing a post-synaptic conductance configuration in close to a sharp transition of the PV+BC ISI), a small decrease in the CCK+BC to PV+BC conductance induces a vast increase in the firing rate of the PV+BCs, even by 20 Hz. This would support the argument that a strong disinhibition of the PV+BCs, mediated by the CCK+BCs can induce a sudden increase in the PV+BC firing rate and mediate the sharp termination of the PYR population. If the FSM dynamics follow those of our grid search of the clamped network model, then a strong inhibition of the CCK+BCs would effectively release the PV+BCs from the CCK+BC inhibition causing a sudden increase in the firing rate of the PV+BC.

### High performance computing simulations

We implement our simulations on Scinet (***Loken et al., 2010***; ***Ponce et al., 2019***) on the Niagara clusters, using 10-12 nodes per simulation with 40 cores per node. Each FSM simulation takes approximately 8 hours real time to be executed. Several FSM simulations were carried out with removal of particular connection. The grid search of post-synaptic weights was also carried out on scinet using 1 node (40 cores per node) and took approximately 3 hours.

### FSM parameter value sets

To rationalize the focus on a subset of the several different cell types in the FSM, we considered the ‘Other Input’ (external drives) received by different cell types (see TABLE 1). From this table, it is clear that AACs receive the smallest amount of other input.

As well, we extracted connectivities between all the different cell types in the FSM. Specifically, we compute an effective weight (EW) which is the connection probability times the synaptic weight. The EWs are shown in TABLES 2 and 3. Note that computed EW values are multiplied by 100 to reduce decimal zeroes.

### Population Rate Model (PRM)

In order to disambiguate the respective role of cell types and their mutual connectivity motifs in the generation and control of theta-gamma coupling in the hippocampus, we developed a reduced, system-level population rate model (PRM). This phenomenological model combines connections and cell types that are identified as critical in generating oscillatory activity and theta-gamma coupling based on previous work (***Chatzikalymniou et al., 2021***) and systematic parameter exploration of the detailed model (***Bezaire et al., 2016b***) here. The purpose of the PRM is the identification and analysis of key mechanisms involved in cross-frequency interactions and oscillatory control over hippocampal microcircuits, following the formalism of population-scale mean-field models.

The PRM describes the time evolution of PYR cell, CCK+BC, PV+BC and BiC mean firing rates (*r*_*PYR*_, *r*_*BiC*_, *r*_*CCK*_ and *r*_*PV*_) as a function of their mutual connectivity motif (FIGURE 3), synaptic weights (*w*_*n*→*m*_, for *n, m* = *P Y R, BiC, CCK* and/or *P V*) membrane rate constants (*α*_*PYR*_, *α*_*BiC*_, *α*_*CCK*_ and *α*_*PV*_), synaptic and axonal delays (*τ*), population-specific inputs (*i*_*PYR*_, *i*_*BiC*_, *i*_*CCK*_ and *i*_*PV*_) as well as intrinsic random fluctuations, and how these collectively contribute to the generation of rhythmic firing rate modulations. The resulting model builds on the well-known ***Wilson and Cowan*** (***1972***) formalism, resulting in the following set of nonlinear delayed stochastic differential equations,

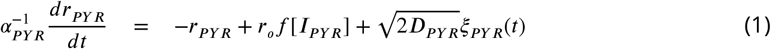

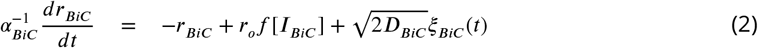

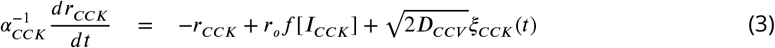

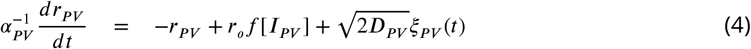

where *r*_*o*_ represents the maximal firing rate of the neurons. Fluctuations in mean firing rates results from the nonlinear integration of presynaptic inputs *I*_*m*_ (here and below, *m* = *P Y R, BiC, CCK* or *P V*) combined to the effect of random perturbations of variance *D*_*m*_ in which *ξ*_*m*_ are uncorrelated zero mean Gaussian white noise process such that < *ξ*_*m*_(*t*)*ξ*_*n*_(*s*) >= *δ*_*nm*_(*t* −*s*), where <> is a temporal average and *δ* is the Dirac delta function. Presynaptic inputs *I*_*m*_ scale firing rates through the firing rate response function

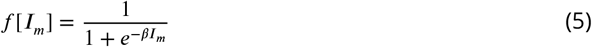

whose gain *β* reflects the steepness of the response. The presynaptic inputs *I*_*PYR*_, *I*_*BiC*_, *I*_*CCK*_ and *I*_*PV*_ are given by

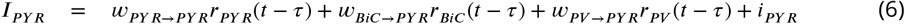

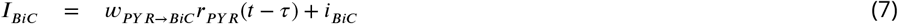

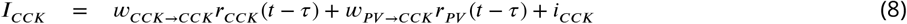

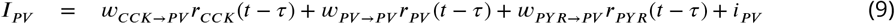

Cell-type specific inputs (*i*_*PYR*_, *i*_*BiC*_, *i*_*CCK*_ and *i*_*PV*_) represent external stimuli modulating the excitability of the different cell populations. A minimal time delay of *τ* = 5*ms* was included to represent finite axonal and synaptic conduction. The parameters *α*_*m*_ represents the membrane rate constant and sets the time scale at which populations respond to stimuli. The coupling constants *w*_*m*→*n*_ represent the synaptic weights from population *m* to population *n* for *n, m* = *P Y R, BiC, CCK* and/or *P V*. Reference parameter values that produce the control output shown in FIGURE 4A are given in TABLE 4.

The PRM exhibits co-modulated oscillations within the gamma and theta frequency ranges. Since the PYR cells constitute the majority of the cells in the network, it is reasonable to consider the PYR cell activity as an LFP proxy. In the absence of multi-compartment representations, various LFP proxies have been used (***Einevoll et al., 2013***) that include number of ‘firing cells’ (***Traub et al., 1992***) which is in essence the PYR cell activity in the PRM. Co-modulated oscillations result from a combination of excitation-inhibition relaxation oscillations and delay-induced oscillations whose peak frequency and power - and resulting dominance - are controlled by a combination of parameters which we explore systematically in FIGURES 5 and 6.

### Simulations and visualizations of the PRM

Each of the thousands of simulations was performed for a duration of 2.0 s with a step size of *dt* = 0.001 s. The differential equations were integrated using a forward Euler method and the output from each of the four different cell type populations in the model was filtered using a band pass filter. For consideration of the theta (*θ*) band frequency, the output was filtered between 3 − 15 Hz while for consideration of the gamma (*γ*) band frequency, the output was filtered between 15 − 100 Hz. The peak frequency was obtained by finding the frequency corresponding to the highest peak in a Welch periodogram (using the *welch* function of the *scipy*.*signal* module in Python with *nperseg* = 1024). The power of the peak frequency was also recorded from the Welch periodogram. Python code to automate the sets of simulations and create all the subsequent visualisations are given in: https://github.com/FKSkinnerLab/PRM.git

#### Heat Maps and Contours

Two-dimensional heat maps of *θ* and *γ* frequencies and power were created for different sets of parameters. For each pair of parameter values that were explored, the range of values for each parameter was divided into 300 equally spaced intervals, creating a 300×300 grid in the two dimensional parameter space. For each point in the grid, the model was simulated using the corresponding values for the parameters being explored while keeping all the other parameters at their reference values. The corresponding frequency and power for the *θ* and *γ* bands were recorded for each point in the grid. Power contours were added to the frequency heat maps to create the plots shown in FIGURES 5 and 6. Six evenly spaced contours were drawn that are representative of the entire range of power values over that grid (using the *MaxNLocator* function of the *matplotlib*.*ticker* module in Python), and were added to the frequency heat map.

#### Power Difference Maps

For comparing the relative power of the *θ* and *γ* oscillations, each of the individual power maps were normalised within their range of values, and the difference of (*θ* − *γ*) normalized powers is calculated. A positive (negative) result indicates a stronger *θ* (*γ*) rhythm due to its larger normalized power.

#### Theta Frequency Plots

For each of the *θ* frequency plots shown in FIGURES 5 and 6, one of the parameters was held constant at a value such that a range of *θ* frequencies over an entire range of values of the other parameter that was being explored had reasonably large power values. The other parameter was varied across the chosen range and the *θ* frequency of the PYR cell was recorded for each point after filtering the output. The ranges and constant values were chosen such that the power of the *θ* oscillations never fell below 10% of its maximum value over the previously explored parameter space for the *i*_*PYR*_ − *w*_*PYR*→*PYR*_ and *w*_*PV*→*CCK*_ − *w*_*CCK*→*PV*_ ranges, and 25% for the *w*_*BiC*→*PYR*_ − *w*_*PYR*→*BiC*_ and *w*_*PV*→*PYR*_ − *w*_*PYR*→*PV*_ ranges.

## Acknowledgements

This research was supported by the Natural Sciences and Engineering Research Council of Canada (NSERC). Computations were performed locally and on the Niagara supercomputer at the SciNet HPC Consortium. SciNet is funded by: the Canada Foundation for Innovation; the Government of Ontario; Ontario Research Fund - Research Excellence; and the University of Toronto.

## Author contributions statement

APC, JL, FKS conceived and designed research; APC, JL, SS, FKS performed research; APC, SS analyzed data; APC, JL, FKS interpreted results; All authors prepared figures and contributed to paper draft writing. All authors reviewed the manuscript.

---

**Figure 1—figure supplement 1.**
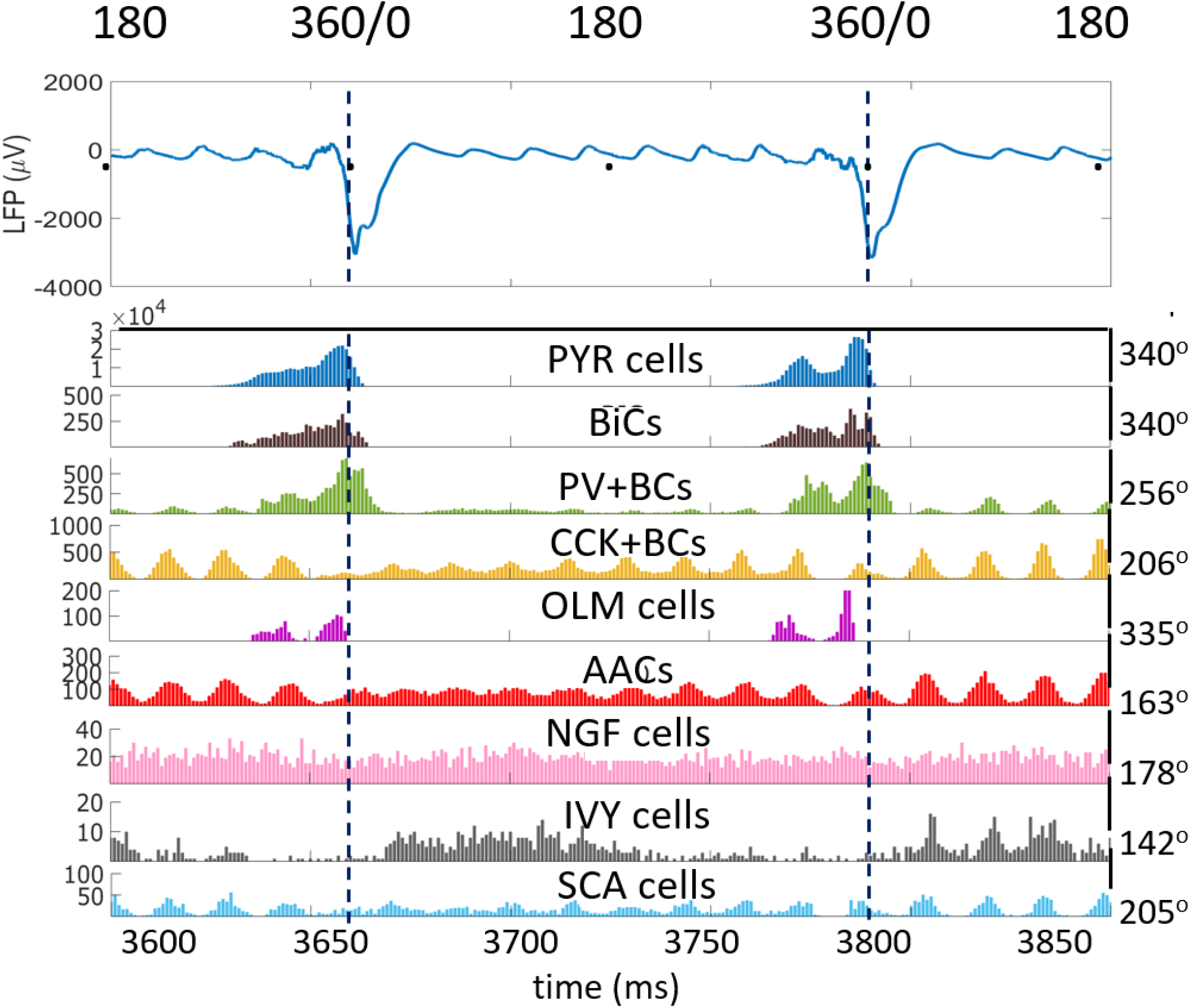
Theta phase preference detail. LFP showing two theta cycles over 300ms with respective histogram plots of the eight inhibitory cell types and the PYR cells showing peak cellular firings relative to the LFP trough.

**Figure 2—figure supplement 1.**
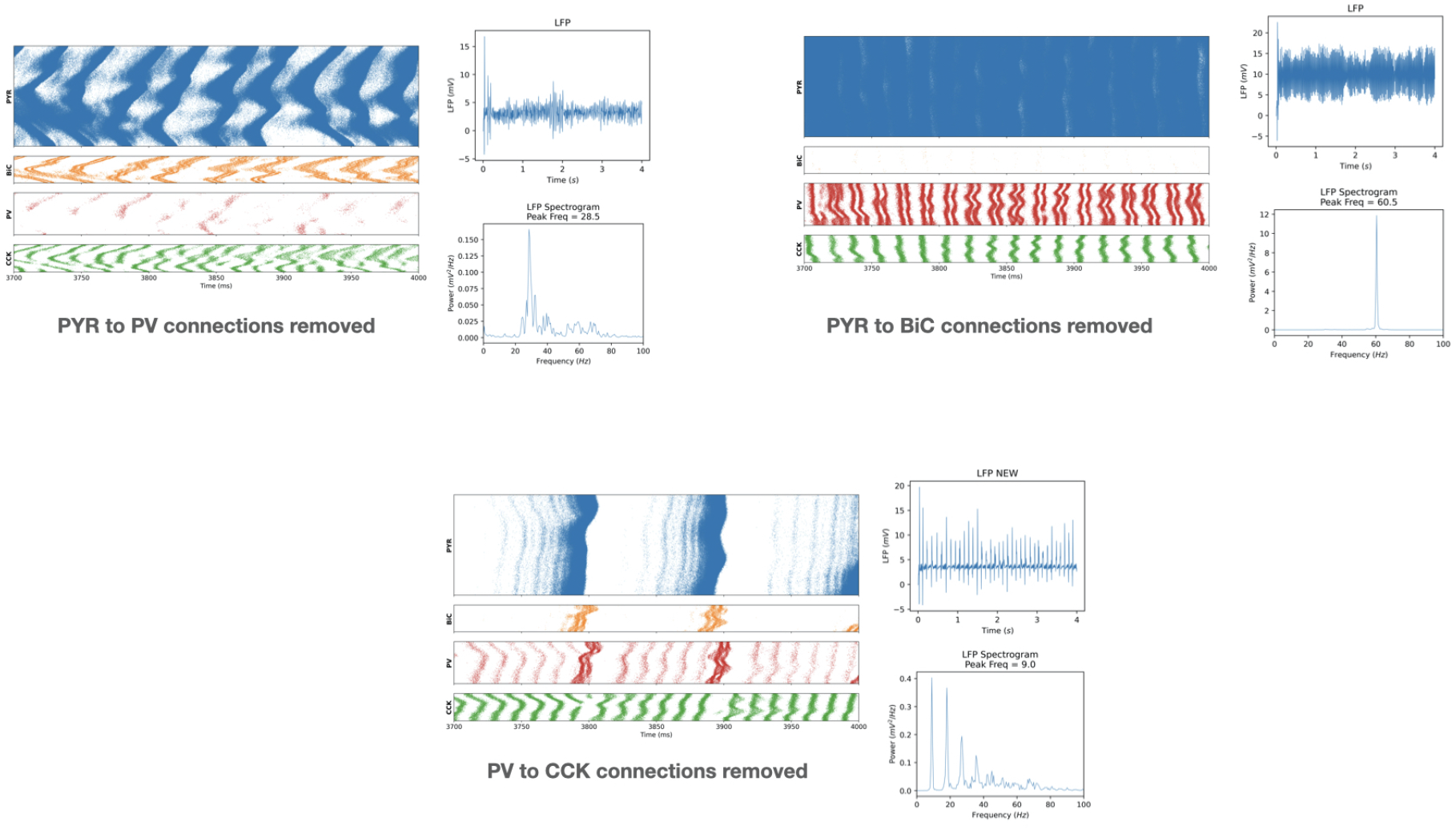
Additional connection removals in the detailed model. **A**. Removing PYR cell to PV+BC connections. (i) Raster plots of cell firings (ii) unfiltered LFP, (iii) Welch’s Periodogram of LFP. **B**. Removing PYR cell to BiC connections. (i) Raster plots of cell firings (ii) unfiltered LFP, (iii) Welch’s Periodogram of LFP. **C**. Removing PV+BC to CCK+BC connections. (i) Raster plots of cell firings (ii) unfiltered LFP, (iii) Welch’s Periodogram of LFP.

**Figure 5—figure supplement 1.**
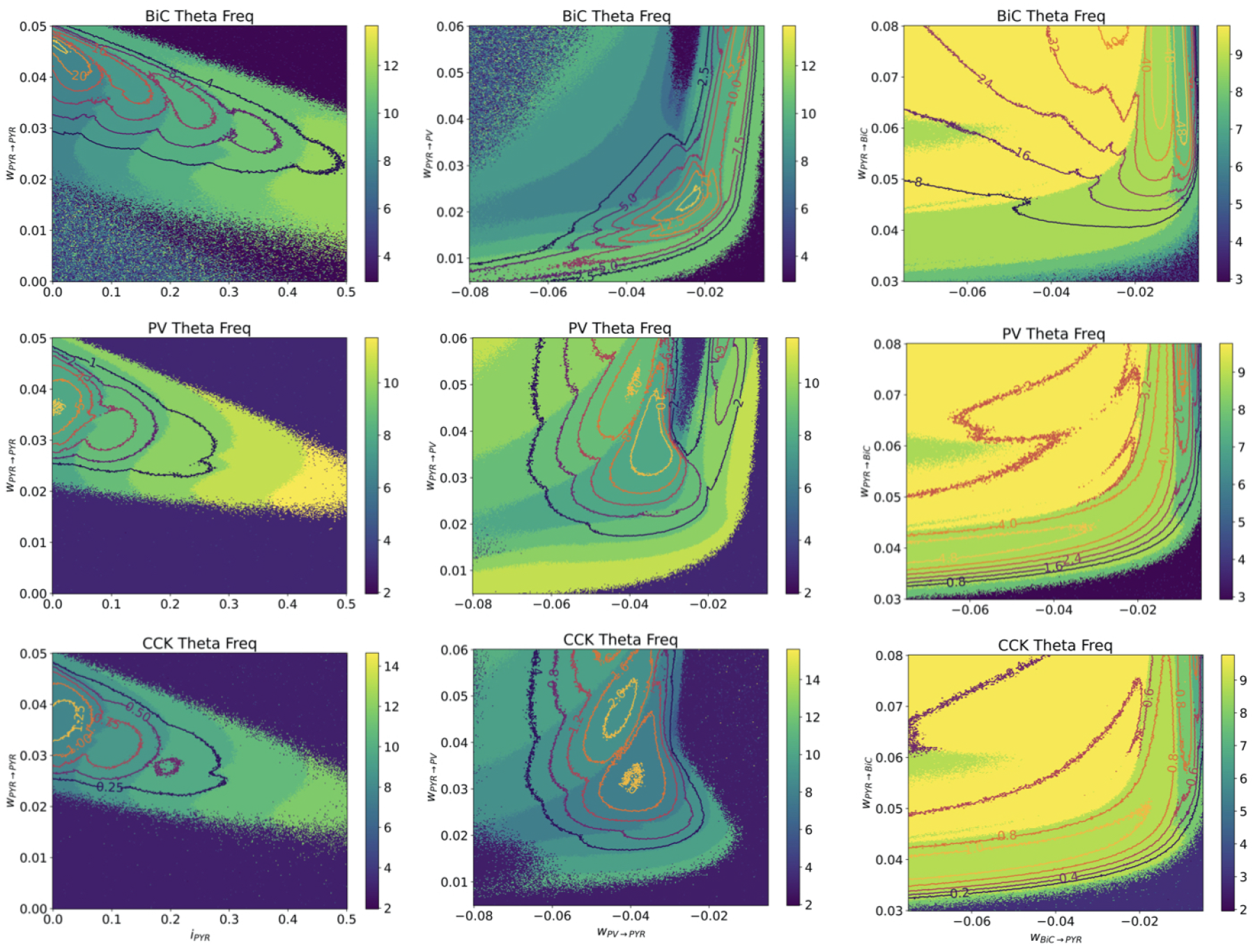
Theta frequency heat maps for other cell types. BiC, PV+BC and CCK+BC cell types of the PRM for the same parameter sets in FIGURE 5 of main text are shown. Only PYR cell theta frequency heat maps are shown in the main text.

**Figure 5—figure supplement 2.**
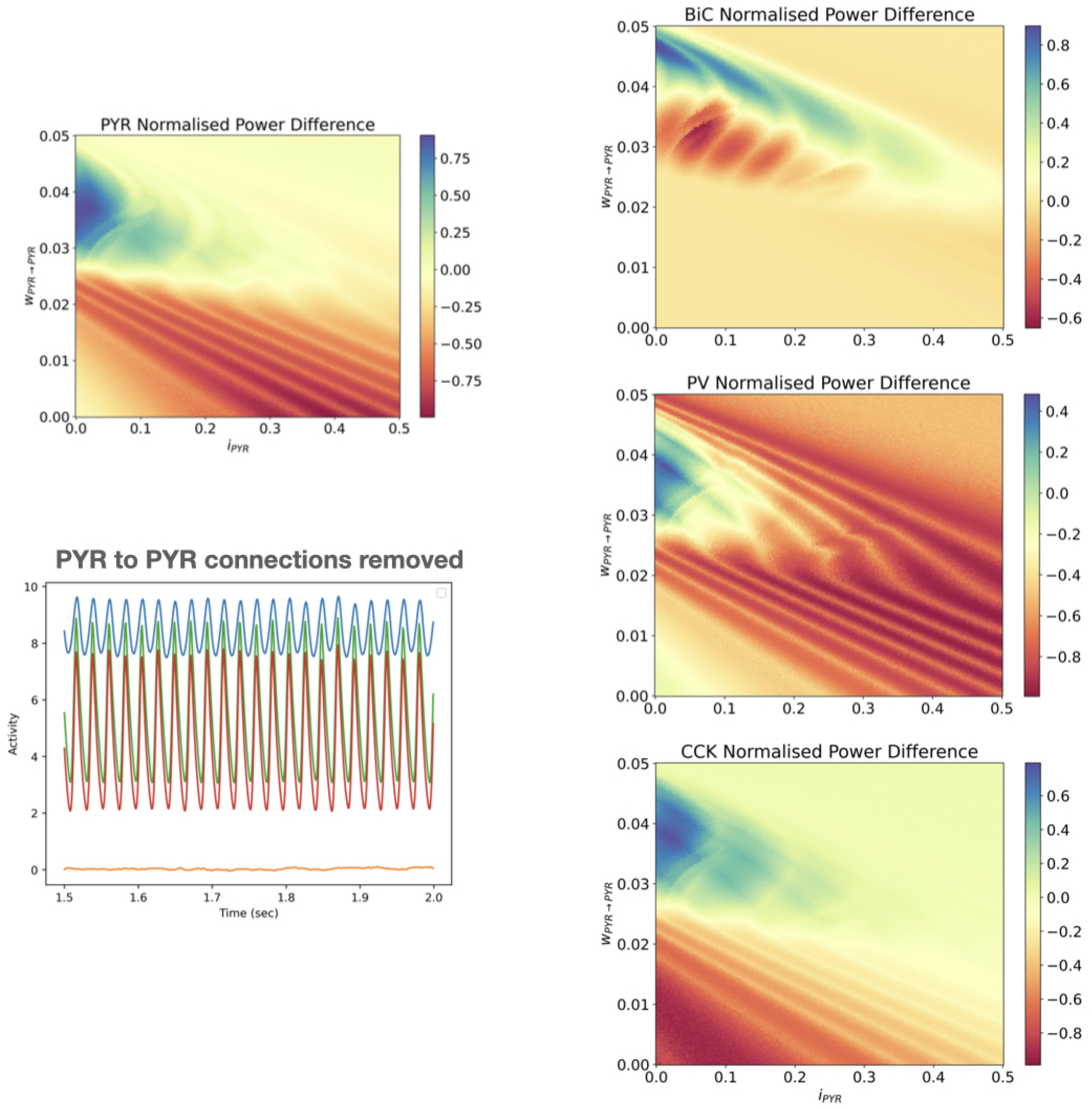
Difference heat maps. Normalized theta and gamma power for *i*_*pyr*_ −*w*_*pyr,pyr*_ are shown for all 4 cell types in difference heat maps (see Methods). Also shown is a plot of the output of cell activities with reference parameter values but when connections between PYR cells are removed (*w*_*pyr,pyr*_ = 0).

**Figure 6—figure supplement 1.**
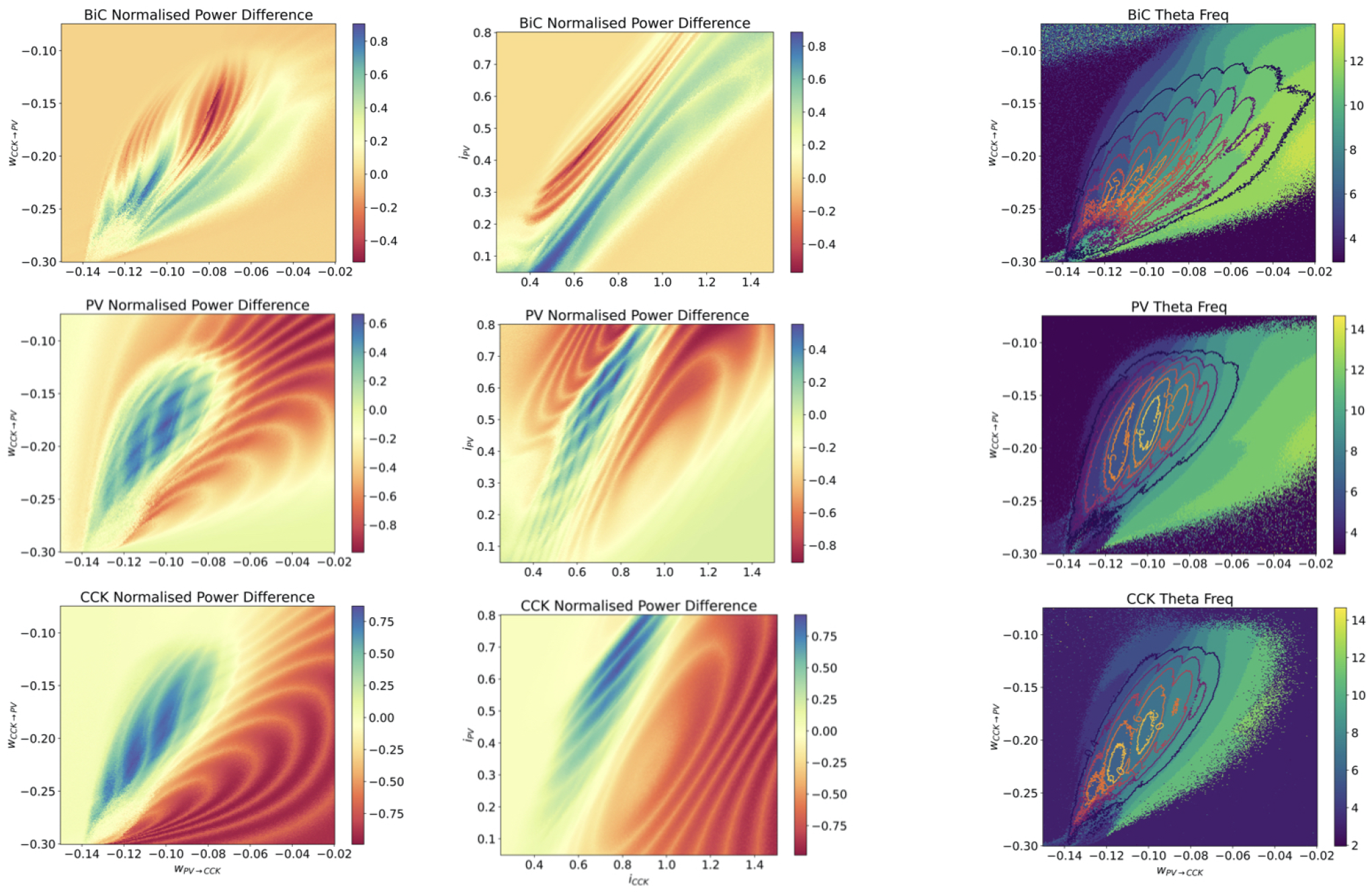
Additional difference heat maps. BiC, PV+BC and CCK+BC cell types of the PRM for the parameter sets in FIGURE 6 of main text are shown. Only PYR cell heat maps are shown in the main text.

**Figure 6—figure supplement 2.**
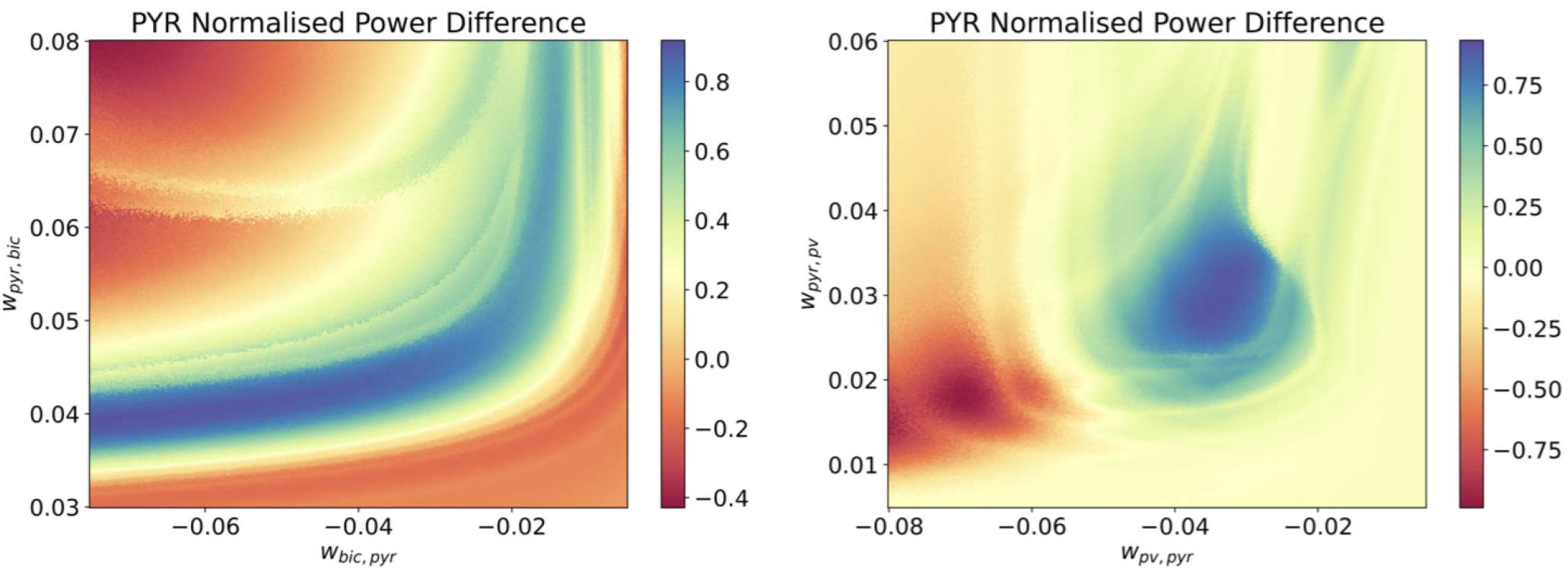
Difference heat maps for PYR-PV+BC and PYR-BiC couplings. Difference heat maps are shown for *w*_*BiC*→*PYR*_ − *w*_*PYR*→*BiC*_ (left) and *w*_*PV*→*PYR*_ − *w*_*PYR*→*PV*_ (right) for PYR cell activity.

