## Supplemental Figures for "Cholecystokinin-expressing (CCK+) basket cells are key controllers of theta-gamma coupled rhythms in the hippocampus" (from another version of this preprint: version 1, 2022-06-03)

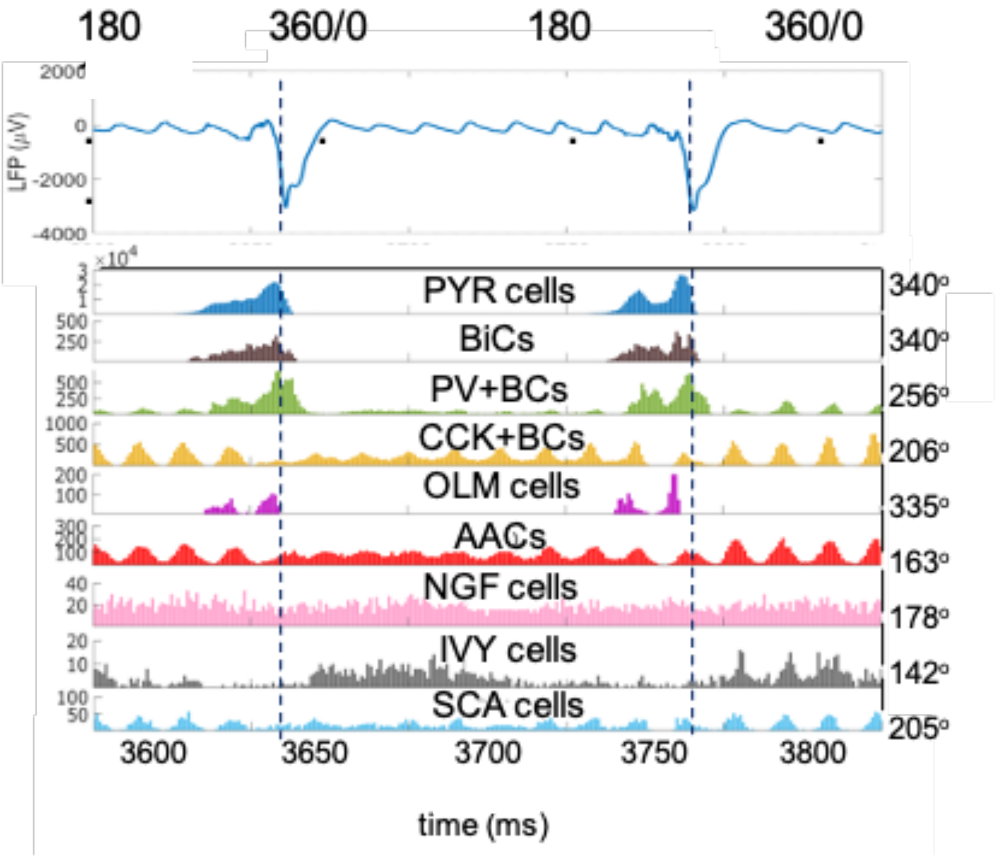
180
360/0
180
360/0
2000
0
-2000
-4000
LFP (μV)
×10^4
PYR cells
340°
BiCs
340°
PV+BCs
256°
CCK+BCs
206°
OLM cells
335°
AACs
163°
NGF cells
178°
IVY cells
142°
SCA cells
205°
3600
3650
3700
3750
3800
time (ms)

**SUPP FIG 1**

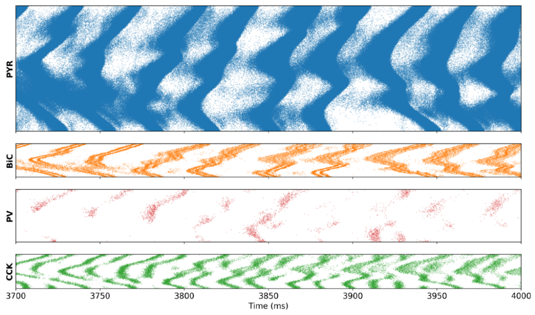


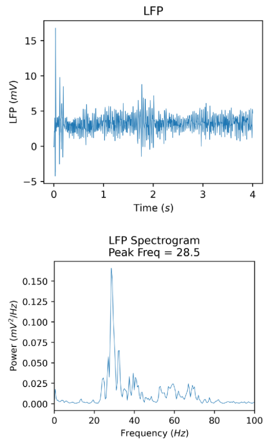


PYR to PV connections removed

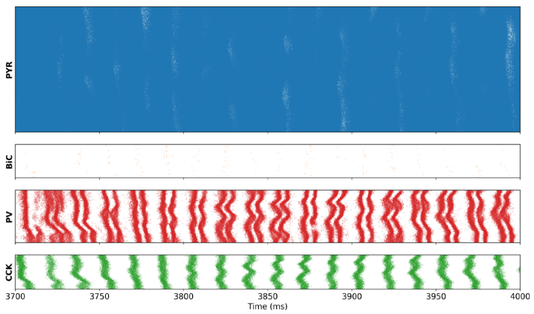


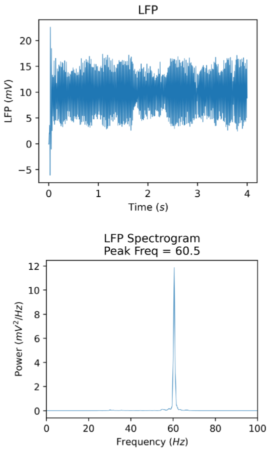


PYR to BiC connections removed

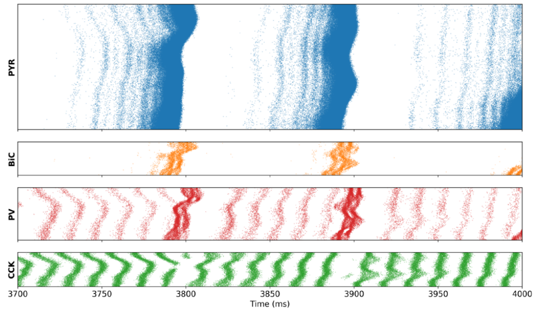


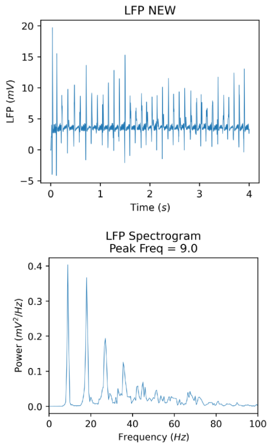


PV to CCK connections removed

**SUPP FIG 2**

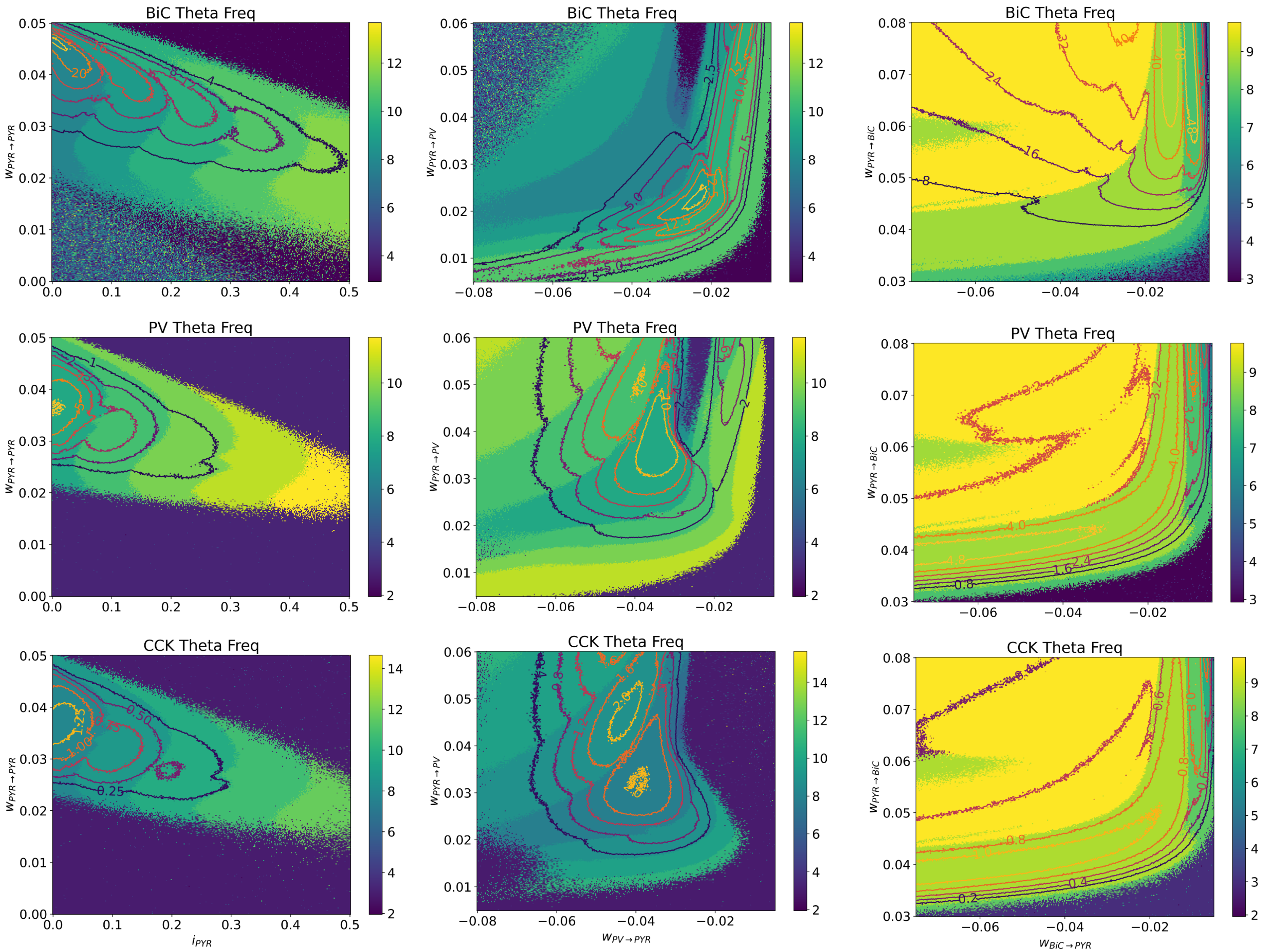

BiC Theta Freq
BiC Theta Freq
BiC Theta Freq
PV Theta Freq
PV Theta Freq
PV Theta Freq
CCK Theta Freq
CCK Theta Freq
CCK Theta Freq
$w_{PYR \to PYR}$
$w_{PYR \to PV}$
$w_{PYR \to BiC}$
$i_{PYR}$
$w_{PV \to PYR}$
$w_{BiC \to PYR}$


**SUPP FIG 3**

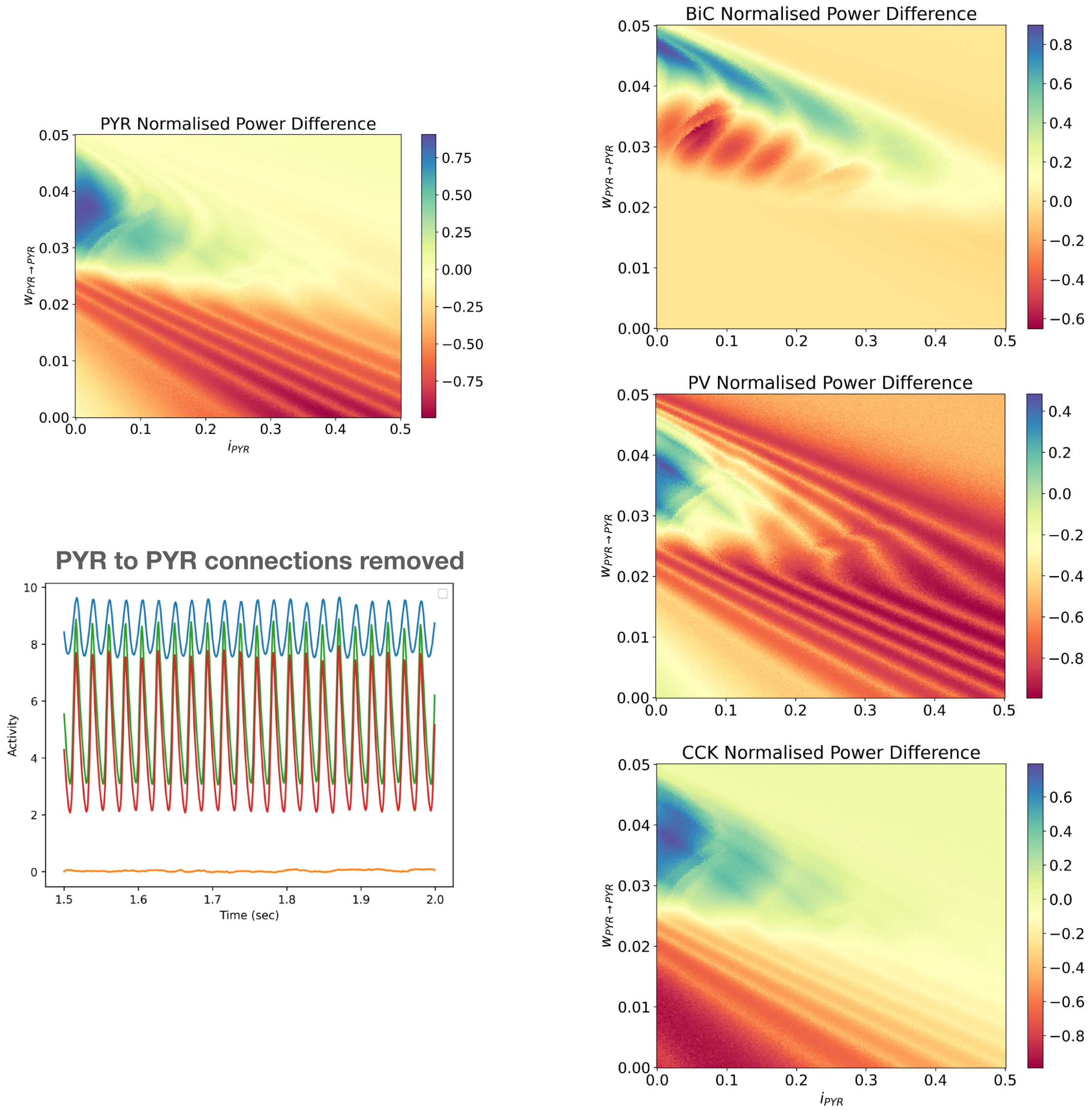
PYR Normalised Power Difference
BiC Normalised Power Difference
PV Normalised Power Difference
CCK Normalised Power Difference
$w_{PYR \to PYR}$
$i_{PYR}$
PYR to PYR connections removed
Activity
Time (sec)

### SUPP FIG 4

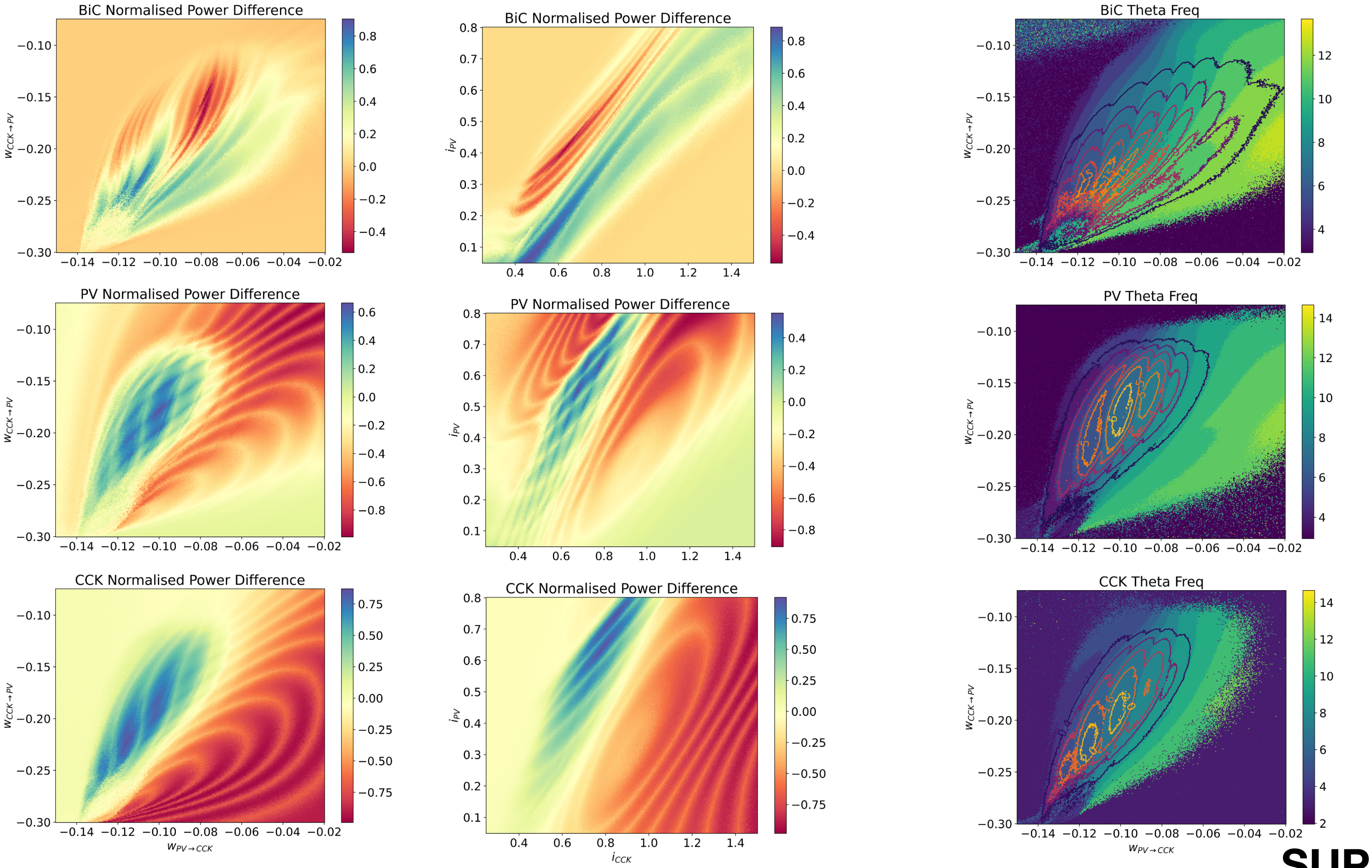
BiC Normalised Power Difference
BiC Normalised Power Difference
BiC Theta Freq
PV Normalised Power Difference
PV Normalised Power Difference
PV Theta Freq
CCK Normalised Power Difference
CCK Normalised Power Difference
CCK Theta Freq
$w_{CCK \to PV}$
$w_{PV \to CCK}$
$i_{PV}$
$i_{CCK}$

**SUPP FIG 5**
